# Deep-learning based 3D segmentation of heterogeneous lizard claw tissue from CT data

**DOI:** 10.64898/2026.07.27.741043

**Authors:** Haleema Sadia, Kate Manges Douglas, Asyiah Bray, Andrea Rummel, Parvez Alam

## Abstract

The accurate segmentation of lizard claws is important as they are materially heterogeneous, comprising both bone and keratinous tissue. This study presents a deep learning framework for the automated segmentation of lizard claw tissues, specifically bone and keratin, from CT imaging data. A dataset comprising 14 lizard claws was used in this work, with annotations generated through a superpixel based labeling approach to provide ground truth reference segmentations. To evaluate the effect of spatial context on segmentation performance, both 2D and 2.5D CNN architectures using DeepLabV3 with ResNet-50, ResNet-101, and Inception-ResNet-v2 backbones were investigated, with predictions subsequently reconstructed into three-dimensional volumes for analysis. Performance was assessed using a leave one out cross validation (LOOCV) strategy and evaluated with 3D Dice Similarity Coefficient (DSC), Intersection over Union (IoU), Sensitivity (Recall), 95th Percentile Hausdorff Distance (HD95), and Relative Volume Error (RVE). Experimental results demonstrate that 2.5D CNN architectures consistently outperform their 2D counterparts across all evaluation metrics, highlighting the importance of incorporating inter-slice contextual information for volumetric tissue segmentation. From amongst the models, the 2.5D Inception-ResNet-v2 achieved the best overall performance, reaching a validation accuracy of 97.5% and producing segmentation results that closely align with ground-truth tissue structures. Our findings demonstrate the effectiveness of 2.5D deep learning approaches for the high accuracy segmentation of heterogeneous lizard claw tissues from CT data, whilst providing a robust framework for automated morphological analysis in comparative anatomical studies.

## 1 Introduction

The locomotion and surface interaction abilities of reptiles have long attracted attention in biomechanics due to their structural efficiency and adaptability [1–7]. In particular, lizard claws play a vital role in essential behaviors such as climbing, gripping, digging, and interacting with a wide range of natural substrates [4–6]. These functions are enabled by their ability to withstand repeated mechanical loading while maintaining structural integrity, which is governed by their complex internal organization and material composition [4]. Unlike homogeneous engineering materials, lizard claws are natural composite structures consisting of a bony core surrounded by a keratinous sheath that contains both predominantly *β*-keratin and hard *α*-keratin homologs, each contributing distinct mechanical functions [8]. The rigid bony core and *β*-keratinous sheath primarily provide resistance to bending, while the presence of hard *α*-keratin enhances energy dissipation and crack resistance [9, 10]. Together, these heterogeneous components interact under mechanical loading to produce a balanced combination of stiffness, toughness, and durability, enabling reliable performance during locomotor activities on rough and irregular substrates [1].

To enable the accurate modeling of lizard claw mechanics, high-resolution CT imaging provides detailed volumetric and radio density information of bone and surrounding composite tissues, enabling comprehensive three-dimensional visualization of internal structure, which can be further leveraged by computational frameworks to model both external morphology and internal features for biomechanical analysis [10–13]. However, despite these advantages, accurately differentiating and modeling the constituent materials from CT data remains challenging because soft keratinous tissues ( *α*-keratin and *β*-keratin) and bone exhibit overlapping CT density distributions and similar grayscale intensities, making simple threshold based segmentation unreliable [8]. Another key challenge is the partial volume effect, which blurs the CT signal at the interface between the surrounding soft tissue and the underlying bone, producing mixed intensities that further obscure the true boundary and reduce segmentation accuracy [14]. Consequently, manual segmentation to distinguish between bone and surrounding composites is often employed; however, it requires expert medical knowledge for accurate annotation and is highly labor intensive, subjective, and time consuming, particularly for large volumetric datasets containing hundreds or thousands of slices [15, 16].

To reduce this burden, traditional histogram based thresholding techniques have been extensively used to differentiate anatomical regions in CT images by exploiting the intensity distribution of imaging data [17] [18]. Traditional machine learning based segmentation methods have been widely applied to CT and MRI image analysis. Among these, SLIC (Simple Linear Iterative Clustering) superpixel segmentation is commonly used to partition images into homogeneous regions based on intensity similarity and spatial proximity, facilitating efficient region based image segmentation [19]. These approaches have been shown to improve boundary adherence and reduce computational complexity compared to pixel wise processing, making them effective for downstream classification and segmentation tasks [20]. In addition, k-means clustering, Gaussian mixture models (GMM), random forests, support vector machines (SVM), and texture-based classifiers, have often been used in combination with intensity based thresholding techniques to improve segmentation performance [13, 21, 22]. By applying statistical learning and handcrafted feature extraction, these methods achieve better accuracy than conventional thresholding alone. However, they remain dependent on manually engineered features and domain-specific preprocessing pipelines, which limit their robustness and adaptability to complex biological structures with heterogeneous morphology [23]. In addition, most methods processed medical image slices independently in two dimensional space without considering contextual continuity along the z-axis. Consequently, these often fail to preserve volumetric consistency and produce discontinuous segmentation boundaries between adjacent slices in 3D CT and MRI datasets [24, 25].

To address the limitations of traditional machine learning approaches, deep learning has emerged as the dominant paradigm for medical image segmentation. Unlike conventional methods that rely on handcrafted features, intensity-based thresholding, or classical classifiers, Convolutional Neural Networks (CNNs) automatically learn hierarchical feature representations directly from imaging data. This enables more robust, adaptive, and accurate characterization of complex anatomical structures across diverse medical imaging modalities [26–28]. Modern frameworks commonly integrate pretrained CNN backbones, such as ResNet-50, ResNet-101, and Inception-ResNet-V2, within architectures like U-Net and DeepLab variants, particularly when training data is limited. Through transfer learning, models first learn general image features such as edges and textures and then adapt them to specific biological structures, resulting in more robust and efficient segmentation performance [29–31]. These models benefit from residual connections, which help improve the training of deeper networks, and further enhances performance by capturing features at multiple scales within CT images [32,33]. These properties are especially useful in biomedical segmentation tasks, where materials such as keratinous tissue and bone exhibit complex and varying morphology across scales.

Although full 3D CNN models such as 3D U-Net and V-Net can achieve high segmentation accuracy by processing entire CT volumes, they are computationally expensive and require significant GPU memory and large annotated datasets [34]. To address this limitation, an intermediate approach known as 2.5D CNN has been proposed. In this method, a small set of consecutive slices (previous, current, and next) is used as input channels [35], allowing the network to capture partial depth information while retaining a 2D convolutional structure. This approach provides an efficient trade-off between 2D and 3D models by preserving inter-slice context with reduced computational cost and memory requirements [36].

Deep learning-based biomedical image segmentation has primarily focused on well-established clinical tasks [26–33, 35]. In contrast, its application to reptilian claw anatomy remains largely underexplored, despite increasing research on reptilian keratin biology and tissue characterization [3–7, 9, 10]. Existing work on reptilian claws has primarily focused on external morphology and surface geometry for biomimetic, robotic, and biomechanical applications [37–40], with limited attention to internal composition. As a result, the segmentation of internal structures such as keratin and bone remains sparsely researched, despite its importance for heterogeneous material characterization and biomechanical modeling. This highlights the need for advanced deep learning-based segmentation methods capable of capturing the internal composite architecture of lizard claws, while concurrently incorporating contextual information for accurate structural representations.

In this study, we propose a 2.5D convolutional neural network (CNN) framework for the segmentation of bone and keratinous tissue in lizard claws, leveraging transfer learning with pretrained deep residual architectures, including ResNet-50, ResNet-101, and Inception-ResNet-v2. The proposed approach captures local volumetric context by integrating information from adjacent CT slices as multi-channel inputs, thereby encoding pseudo-3D spatial continuity while retaining the computational efficiency of 2D convolutions. To assess the impact of contextual information, the 2.5D model is compared against a conventional 2D CNN baseline that processes each slice independently without incorporating inter-slice dependencies. This comparative analysis demonstrates the role of volumetric context in enhancing segmentation performance, particularly for structurally similar tissues such as bone and keratin. The training labeled data for the proposed network is generated through a hierarchical annotation strategy combining SLIC-based superpixel segmentation, intensity driven coarse classification, and histogram valley guided intra-class superpixel merging. Unlike conventional fixed threshold segmentation, which is sensitive to intensity overlap, noise, and inter scan variability, this approach avoids rigid cutoffs by incorporating spatial context and distribution aware refinement, resulting in more robust delineation of heterogeneous tissues with overlapping CT intensity profiles [19] [20] [18] [41]. The proposed study is intended to support downstream biomechanical analysis and finite element modeling by providing more realistic representations of heterogeneous tissue distributions within the anatomy of lizard claws from a variety of different species.

## 2 Research Methodology

### 2.1 Dataset Preparation

#### 2.1.1 Lizard Dataset

In this study total of fourteen lizard claws CT datasets are used as shown in Figure 1. The datasets included both forelimb and hindlimb claws from multiple lizard species. Datasets consisted of *Basiliscus vittatus* (hind), *Basiliscus vittatus* (fore), *Cophosaurus texanus*, *Cordylus giganteus* (hind), *Heloderma suspectum*, *Phrynosoma modestum* (fore), *Salvator merianae* (hind), *Sceloporus arenicolus* (hind), *Sceloporus arenicolus* (fore), *Crotaphytus collaris* (hind), *Crotaphytus collaris* (fore), *Uma notata* (hind), *Varanus salvator* (fore), and *Varanus salvator* (hind). Each claw dataset comprised approximately 1000 sequential DICOM slices obtained from high resolution CT imaging. The dataset was utilized in all three anatomical planes, namely axial, coronal, and sagittal views of each claw, to enrich the spatial representation of the structures.

**Figure 1:**
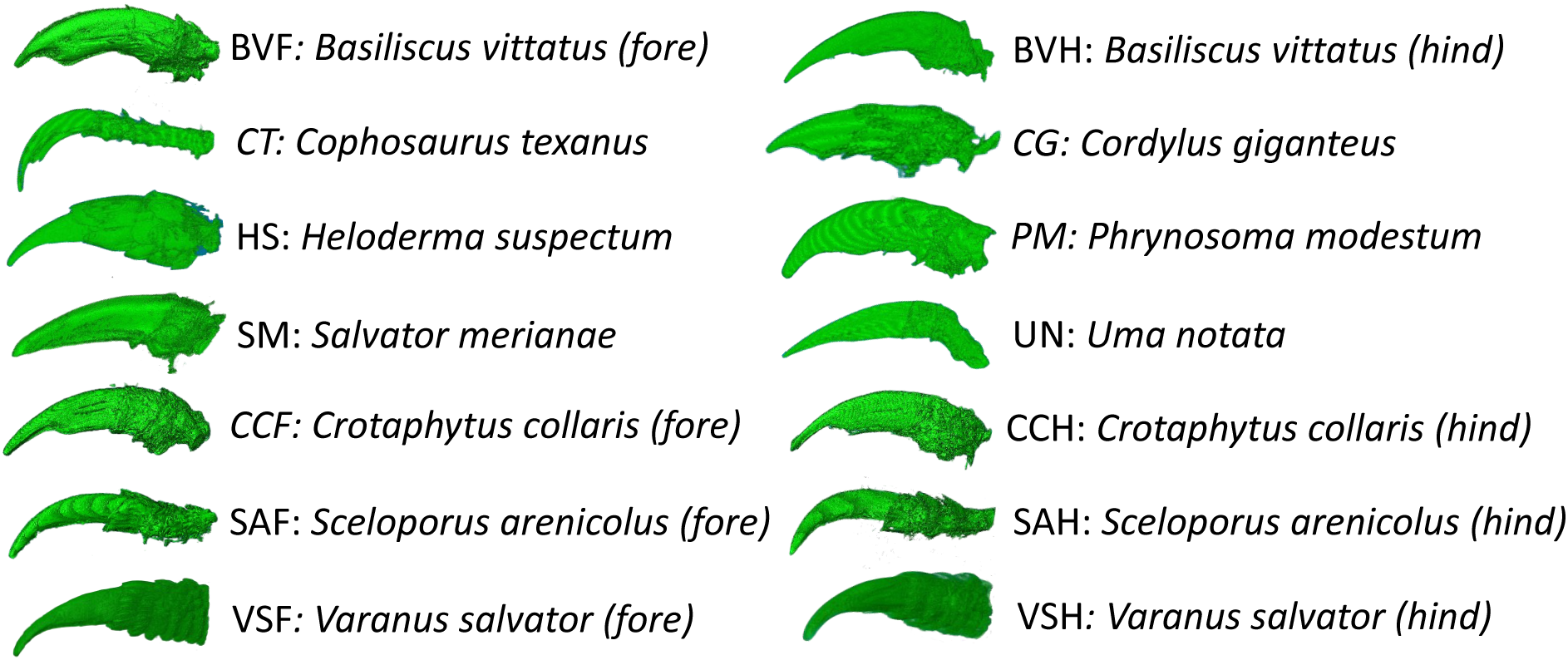
Overview of the CT dataset used in this study. The dataset consists of claw scans from multiple lizard species, including *B. vittatus* (hind and fore), *C. texanus*, *C. giganteus* (hind), *H. suspectum*, *P. modestum* (fore), *S. merianae* (hind), , *U. notata* (hind), *C. collaris* (hind and fore), *S. arenicolus* (hind and fore), and *V. salvator* (fore and hind). Each claw volume consists of 1000 consecutive DICOM CT slices, providing high-resolution volumetric data for detailed analysis of bone and keratinous tissues.

#### 2.1.2 Conversion of CT Hounsfield Units to Normalized Grayscale Intensities (0–255)

CT imaging produces cross sectional representations of internal structures by quantifying X-ray attenuation across tissues, which are subsequently expressed as grayscale intensities. The Hounsfield Unit (HU) scale provides a standardized quantitative measure of tissue radiodensity in computed tomography [42]. On this scale, air is assigned a value of approximately −1000 HU, water is defined as 0 HU, while bone structures typically range from about +1000 HU for cancellous bone to +2000 HU for dense cortical bone. Owing to this high attenuation, bone structures are readily distinguishable from surrounding soft tissues in CT images [42–44].

Although precise HU ranges for reptilian keratin are not well established in literature. Bone tissue, composed mainly of hydroxyapatite and collagen, is significantly more mineralized than keratinous structures, which are protein-based and minimally mineralized [45,46]. As a result, bone exhibits higher HU values due to greater X-ray attenuation, while keratinous regions correspond to lower HU ranges. Based on the literature, we define the threshold for background (black) as −1000 to 0, for bone as values above 1000, and for the keratinous tissue as those with values between 0 and 1000. DICOM images were converted to HU using scanner-specific calibration parameters, namely the Rescale slope and Rescale intercept provided in each file. The resulting HU images were then normalized to an 8-bit grayscale range (0–255) to standardize intensity values for further analysis. This normalization allowed consistent comparison across samples. Based on the defined thresholds, the images were subsequently segmented into air, keratinous tissue, and bone, as summarized in Table 1.

**Table 1:** Predefined 8-bit grayscale intensity thresholds used for the classification of air, keratinous tissue, and bone regions in lizard claw CT images.

| **Tissue Type** | **Normalized Intensity (0–255)** |
| --- | --- |
| Air | 0 to 20 |
| Keratin | > 20 to 200 |
| Bone | > 200 |

#### 2.1.3 Superpixel-Based Annotation Framework for CT Claw Slices

To automate the annotation of CT claw slices, we propose a superpixel-based segmentation framework as a means of generating labeled training data for three tissue classes as shown in Figure 2, namely air, keratin, and bone. The proposed hierarchical annotation strategy integrates SLIC (Simple Linear Iterative Clustering) based superpixel segmentation [41], intensity driven coarse classification, and histogram valley guided intra-class superpixel merging [18] to generate anatomically consistent and noise robust labels while preserving spatial and structural boundaries in CT claw images. SLIC is a superpixel algorithm that groups neighboring pixels into compact, perceptually homogeneous regions based on both intensity similarity and spatial proximity, thereby reducing image complexity while preserving meaningful boundaries [19]. Histogram valley guided merging refers to the use of minima (valleys) in the intensity histogram, which typically indicate natural separations between tissue classes, to refine and merge initially over-segmented regions into more anatomically coherent structures. Our approach consists of two primary phases:

**Figure 2:**
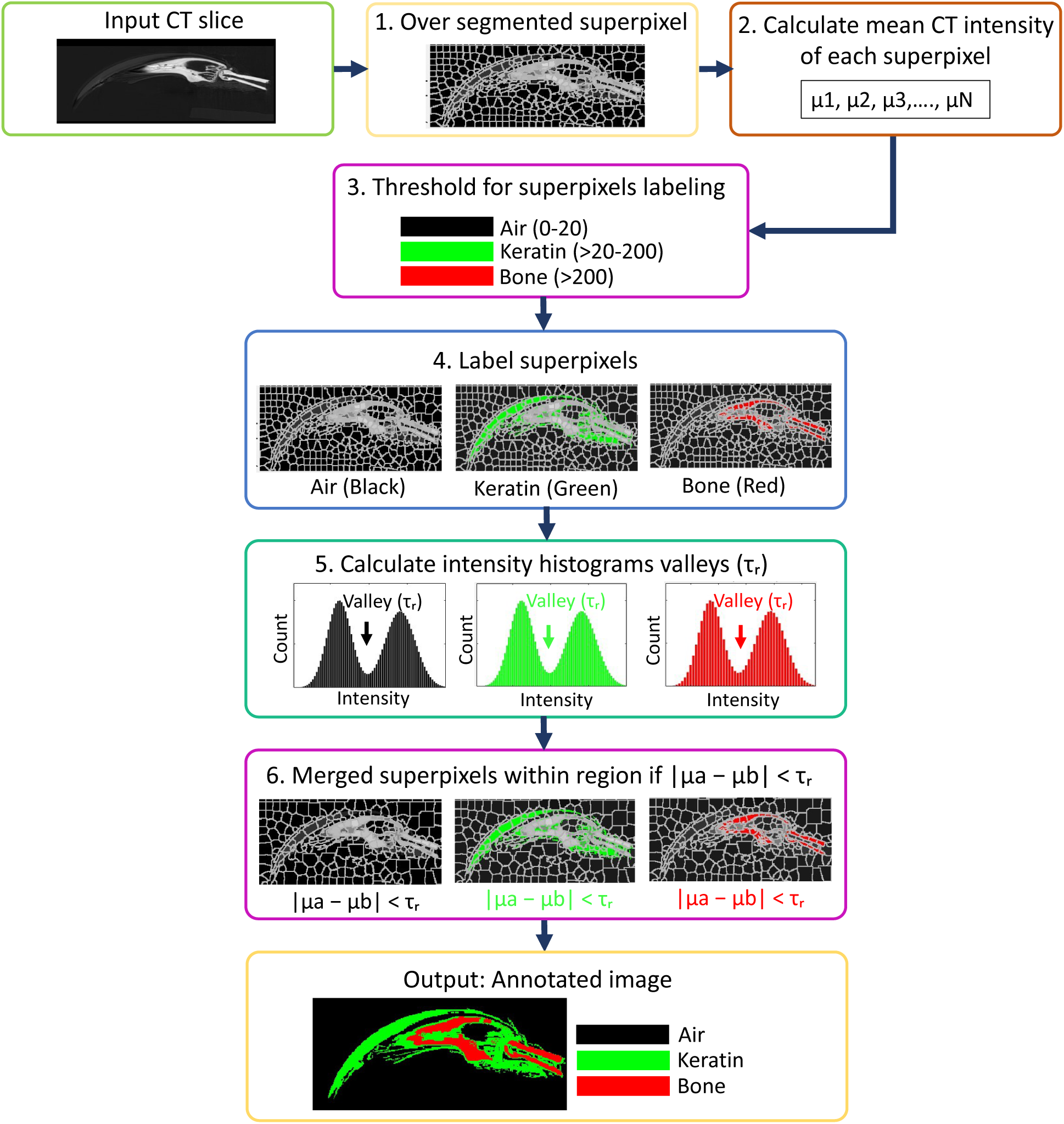
Workflow of the proposed automatic data-driven superpixel merging framework for CT-based claw tissue annotation. The input CT slice (256 × 256 pixels) is initially over-segmented into 2000 superpixels to preserve fine local structural details [47]. The mean intensity of each superpixel is then computed and used for initial labeling based on predefined intensity thresholds corresponding to air, keratin, and bone regions. Subsequently, intensity histograms are constructed within each labeled region, and valley points are automatically detected to derive data-driven thresholds. These valley-based thresholds are then used to iteratively merge adjacent superpixels with similar intensity characteristics, resulting in the final refined segmentation of air, keratin, and bone tissues.

##### 1. Initialization of superpixels

First, the input CT image is preprocessed by normalizing the Hounsfield Unit (HU) intensity values to a 0–255 grayscale range (as shown in Table 1) resulting in the image, Equation 1:

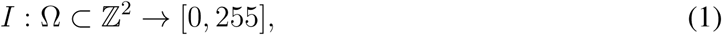

where *I*(*x, y*) represents the normalized intensity at pixel location (*x, y*), and Ω denotes the image domain. Each pixel is represented by a feature vector, Equation 2:

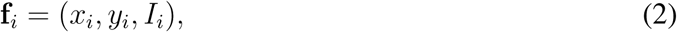

where (*x_i_, y_i_*) are spatial coordinates and *I_i_* is the normalized intensity. Subsequently, an oversegmentation step is performed to generate an initial set of superpixels. These superpixels serve as the basic processing units for subsequent analysis. The image is partitioned into *K* superpixels using the SLIC algorithm. The grid interval is defined as, Equation 3:

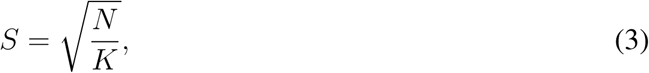

where *N* is the total number of pixels. Each superpixel is initialized with a cluster center, Equation 4:

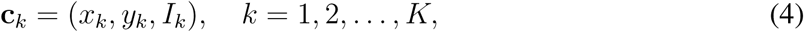

where (*x_k_, y_k_*) and *I_k_* denote the spatial location and intensity of the cluster center. The spatial and intensity distances between pixel *i* and cluster center *k* are defined as in Equations 5 and 6:

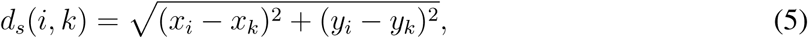

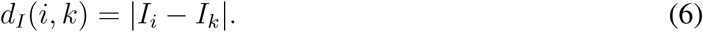

These are combined into the SLIC distance metric, Equation 7:

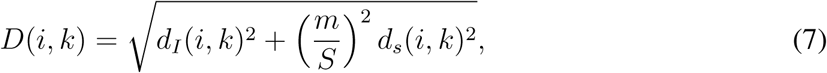

where *m* is the compactness parameter controlling the trade-off between spatial proximity and intensity similarity. Each pixel is assigned to the nearest cluster, Equation 8:

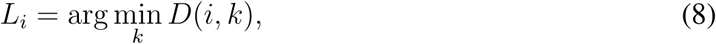

where *L_i_* is the label of pixel *i*. After assignment, cluster centers are updated iteratively as shown in Equation 9:

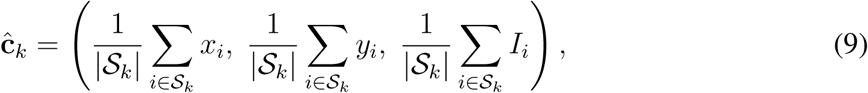

where *S_k_* is the set of pixels belonging to superpixel *k*. Each superpixel is then represented by its mean intensity, Equation 10:

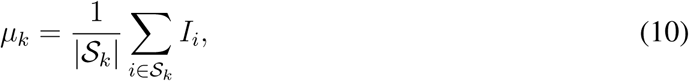

Based on predefined intensity intervals derived from Table 1, each superpixel is assigned to a tissue class, Equation 11:

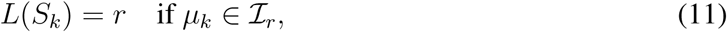

where *r* ∈ {air, keratin, bone} and *I_r_* denotes the intensity range of class *r*. This provides a coarse anatomical labeling of the image.

##### 2. Histogram-Based Superpixel Merging

Following the initial over-segmentation and coarse labeling, histogram analysis is performed within each tissue class. For each class *r*, a histogram of superpixel intensities is constructed, and superpixels with similar intensity distributions are merged to obtain spatially coherent regions, Equation 12:

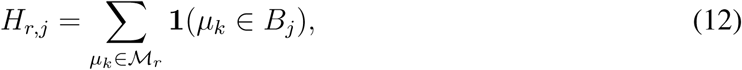

where *H_r,j_* denotes the histogram count for tissue class *r* in the *j*-th intensity bin. The variable *r* ∈ {air, keratin, bone} indexes the tissue class under consideration, while *j* indexes the histogram bins. *M_r_* represents the set of mean intensity values *µ_k_* of all superpixels assigned to class *r*. Each *µ_k_* corresponds to the average intensity of the *k*-th superpixel. *B_j_* denotes the *j*-th predefined intensity bin, which partitions the intensity range into discrete intervals. The function **1**(·) is an indicator function, which takes the value 1 if the condition inside the parentheses is true (i.e., *µ_k_* ∈ *B_j_*), and 0 otherwise. Thus, the equation counts how many superpixels assigned to class *r* have mean intensities falling within bin *B_j_*, forming the histogram *H_r,j_*. Dominant intensity modes are identified as histogram peaks, Equation 13:

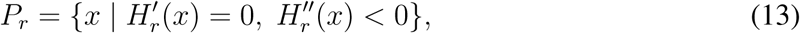

where the condition 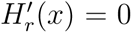 identifies stationary points of the histogram (i.e., locations where the slope is zero), while 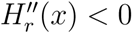 ensures that these stationary points correspond to local maxima. Therefore, *P_r_* consists of all intensity values *x* that satisfy both conditions, representing the peak intensity modes of the histogram for class *r*. Histogram valleys are defined as shown in Equation 14:

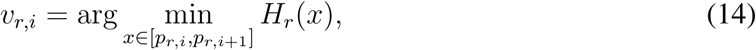

forming the set *V_r_* = {*v_r,_*_1_*, v_r,_*_2_, … }. Here, *v_r,i_* denotes the *i*-th valley of the histogram corresponding to tissue class *r*. The interval [*p_r,i_, p_r,i_*_+1_] defines the intensity range between two consecutive histogram peaks (modes). The operator arg min identifies the intensity value *x* within this interval at which the histogram *H_r_*(*x*) attains its minimum value, corresponding to a local valley separating two adjacent peaks. An adaptive merging threshold is computed as shown in Equation 15:

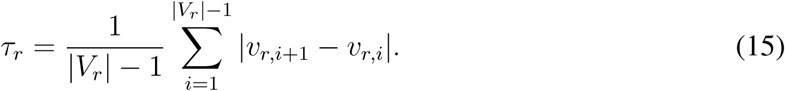

Finally, spatially adjacent superpixels *S_a_* and *S_b_* within the same class are merged if Equation 16 is satisfied:

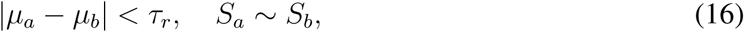

where *S_a_* ∼ *S_b_* denotes spatial adjacency. This merging strategy combines only intensity-consistent and spatially connected superpixels, producing anatomically coherent and noise-robust annotation masks. The superpixel-based annotated dataset provides region-wise labels for air, keratin, and bone, used to train the CNN models to learn discriminative features from each anatomical region for accurate segmentation of lizard claw CT data.

#### 2.1.4 2.5D CNN Architecture

In this work, we implement a 2.5D convolutional neural network (CNN) framework for CT image segmentation as shown in Figure 3. CT images are inherently three-dimensional, and anatomical structures such as air, keratin, and bone extend continuously across neighboring slices. A 2.5D input representation was constructed by stacking three consecutive CT slices as separate channels, where the central slice represents the target image and the adjacent slices provide contextual information from preceding and succeeding anatomical locations. A stack of three consecutive slices is used to provide local contextual information while maintaining a balance between spatial continuity and computational efficiency; larger stacks (e.g., 5 or 7 slices) were avoided to prevent redundant contextual overlap and increased computational complexity [48]. This strategy allows the network to capture local volumetric continuity and inter-slice anatomical relationships while retaining the simplicity of 2D convolutions. For each target slice *d_i_*, a stack of three consecutive CT slices is constructed as Equation 17:

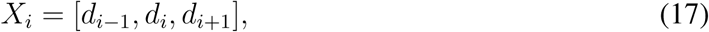

where *d_i_*_−1_, *d_i_*, and *d_i_*_+1_ denote the previous, current, and subsequent CT slices, respectively. The resulting input tensor is represented as in Equation 18:

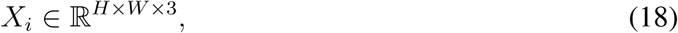

where the first channel corresponds to the previous slice, the second channel corresponds to the target slice, and the third channel corresponds to the next slice. *H* and *W* denote the image height and width. The central slice *d_i_* serves as the prediction target, while neighboring slices provide additional anatomical context. The stacked input tensor *X_i_* is passed through a DeepLabV3 architecture, where three different backbone networks ResNet50, ResNet101, and Inception-ResNetV2 are employed as transfer learning feature extractors.

Each backbone is used to learn hierarchical feature representations from the input data by Equation 19:

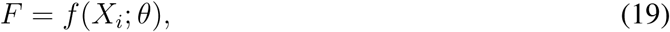

where *f* (·) denotes the backbone network, and *θ* represents the set of all learnable parameters of the backbone network.

A key component of the proposed framework is convolution operations, which enable the network to learn meaningful anatomical features from stacked CT slices (*X_i_*). Convolutional kernels are applied across both the spatial dimensions and the slice-channel dimension of the input tensor. For an input tensor (*X_i_*), the convolution operation produces an output feature map (O), Equation 20:

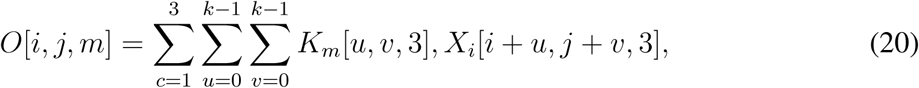

where *O*[*i, j, m*] denotes the output feature value at spatial location (*i, j*) in the *m*-th output feature map, *K_m_* denotes the *m*-th convolutional kernel. The indices *u* and *v* correspond to the spatial coordinates of the convolutional kernel of size *k* × *k*. Through this operation, the network learns spatial features within each slice while simultaneously exploiting contextual information from neighboring slices. The convolutional kernels are applied with a stride *s*, which determines the step size of the kernel movement across the input tensor. To preserve spatial information near image boundaries, zero-padding of size *p* may be applied to the input. Given a kernel size *k*, the spatial dimensions of the resulting feature maps are computed by Equations 21 and 22:

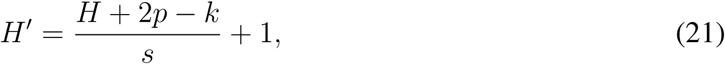

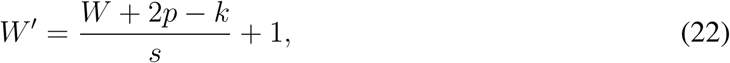

where *H* and *W* denote the height and width of the input feature map, respectively, and *H*^′^ and *W* ^′^ denote the spatial dimensions of the output feature maps. Although the input to the network is a 2.5D tensor, the spatial convolution is performed independently over the height and width dimensions, while the channel dimension is aggregated within the convolutional operation.

**Figure 3:**
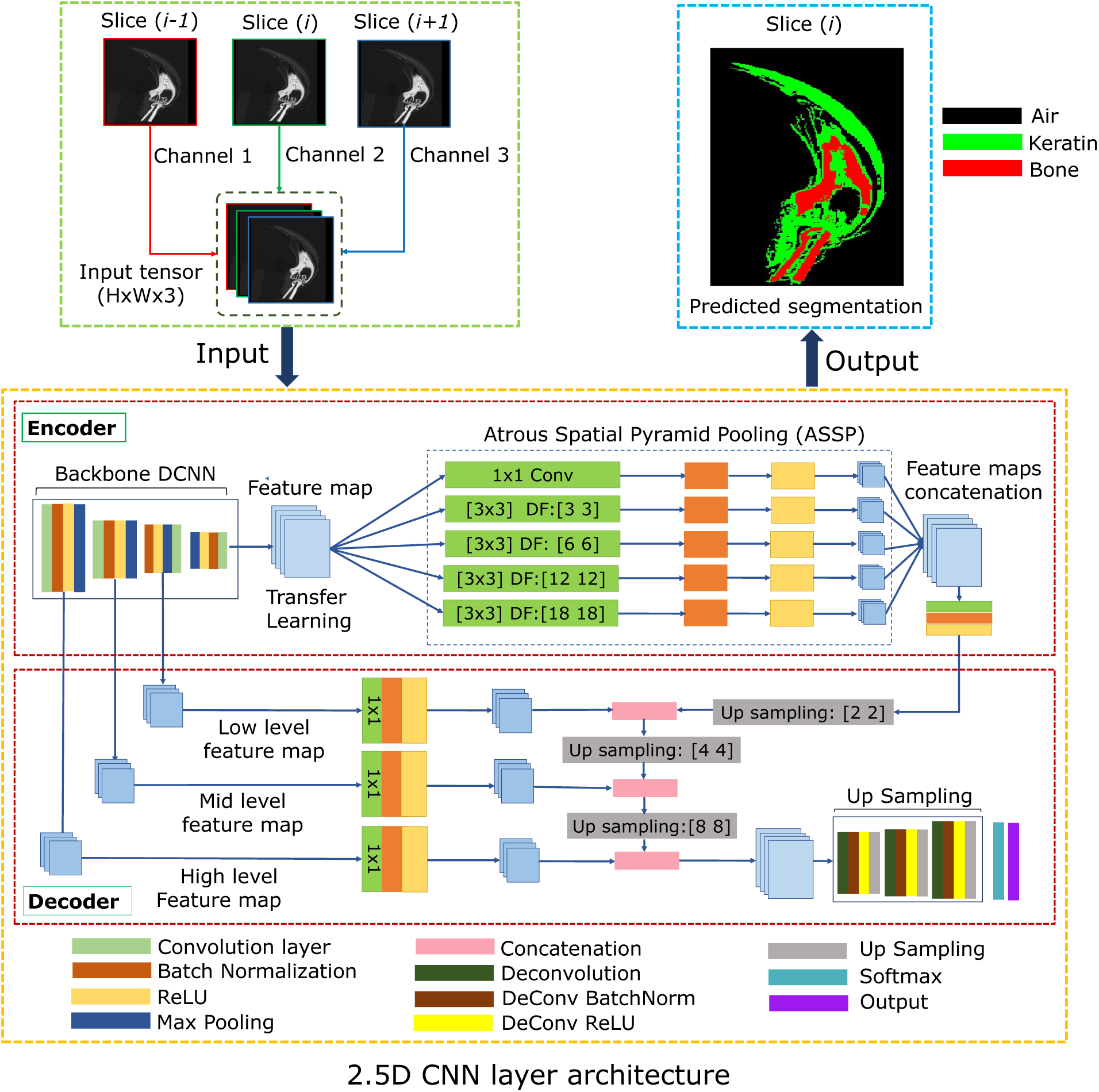
Detailed illustration of the network architecture comprising an encoder-decoder structure. The encoder extracts features from the input data through multiple layers of convolutional operations, progressively reducing spatial dimensions while retaining important information. After feature extraction, an ASSP (Atrous Spatial Pyramid Pooling) module is applied to capture multi-scale contextual information. These concatenated feature maps are then passed through an upsampler before being sent for further upsampling and refinement in the decoder for accurate predictions and pixel classification

Following convolution, a Rectified Linear Unit (ReLU) activation function is applied elementwise to introduce non-linearity and enhance the representational capacity of the network. The ReLU activation is defined as in Equation 23:

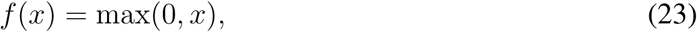

where *x* represents an element of the convolutional feature map prior to activation. This operation suppresses negative responses while preserving positive feature activations, thereby facilitating efficient gradient propagation during training.

Subsequently, max pooling is employed to reduce the spatial dimensions of the feature maps while retaining the most discriminative features. Max pooling partitions each feature map into non-overlapping regions of size *p* × *p* and selects the maximum value within each region. For a feature map *F*, the pooled output *P* is given by Equation 24:

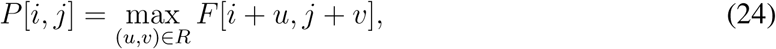

where *R* denotes the pooling region. This operation reduces computational complexity, increases robustness to small spatial variations, and preserves salient anatomical features extracted from the multi-slice CT input.

To stabilize the training process and improve convergence, batch normalization is applied after convolutional layers. Batch normalization standardizes activations within each mini-batch to have zero mean and unit variance. For a batch of activations *x*, the normalized output *x*^ is given by Equation 25:

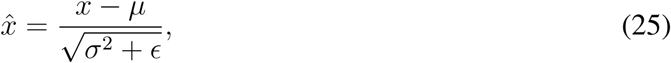

where *µ* and *σ*^2^ denote the mean and variance computed over the mini-batch, and *ɛ* is a small constant added for numerical stability. The normalized activations are then rescaled and shifted using learnable parameters *γ* and *β*, yielding the final output, Equation 26:

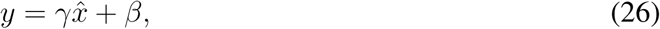

where *y* represents the transformed activation passed to the subsequent layer.

After feature extraction by the backbone network, the resulting feature map is forwarded to the Atrous Spatial Pyramid Pooling (ASPP) module as shown in Figure 3, which enhances multi-scale contextual representation. The ASPP applies parallel atrous (dilated) convolutions with different dilation rates (3, 6, 12, and 18), together with a 1 × 1 convolution branch. This encoder output is then passed to the decoder for final pixel-wise segmentation of anatomical structures in the 2.5D CT input.

The decoder progressively restores the spatial resolution of the encoded feature maps while integrating multi-level contextual information extracted from the backbone and the Atrous Spatial Pyramid Pooling (ASPP) module. Since the input consists of a 2.5D tensor *X_i_* ∈ R*^H^*^×^*^W^* ^×3^, the decoder operates on 2D feature maps with enriched inter slice contextual representations embedded in the channel dimension. The ASPP output is first upsampled by a factor of 2 and fused with corresponding high-level encoder features via skip connections to recover spatial structure. This process is repeated with intermediate and low-level encoder features through successive 2 upsampling steps, enabling hierarchical integration of semantic information with fine-grained spatial details such as edges and textures. Finally, the fused feature map is passed through a 1 × 1 convolution to generate class-wise score maps, followed by a pixel-wise softmax to obtain the final segmentation probabilities. As the model is based on a 2.5D CNN, all decoder operations remain 2D, while inter-slice context is implicitly encoded in the learned features. The framework addresses the limitation of 2D approaches that analyze slices independently, which may lose important spatial context from the inherently three-dimensional nature of CT data [36].

To train the semantic segmentation model, key hyperparameters were systematically tuned for optimal performance. Different optimizers, including Adam, SGD, and SGD with momentum, were evaluated along with learning rates of 0.0001, 0.001, and 0.01. Batch sizes of 16, 32, and 64 were tested to balance computational efficiency and stability. The model was trained for up to 100 epochs with early stopping to prevent overfitting, and a learning rate scheduler was used to gradually reduce the learning rate during training. Categorical cross-entropy was employed as the loss function, and dropout (0.3–0.5) was applied for regularization. Hyperparameters were iteratively refined using the validation set to achieve optimal and stable performance.

#### 2.1.5 Data Augmentation

To improve the robustness and generalization of the segmentation model, several data augmentation techniques were applied to the CT-based lizard claw dataset during training, include horizontal and vertical flipping, small rotations within ±10^◦^, and random cropping of 10%–50% of the image area to improve spatial invariance and localization ability. In addition, zero-mean Gaussian noise with *σ* = 0.01–0.05 was added to simulate CT imaging artifacts and enhance robustness to intensity variations and acquisition noise [49]. The dataset was utilized in all three anatomical planes, namely axial, coronal, and sagittal views of each claw, to enrich the spatial representation of the structures.

### 2.2 Validation Metrics

The performance of the proposed 2.5D CNN for lizard claw CT image segmentation was quantitatively evaluated using several widely adopted metrics in medical image analysis. These metrics assess the agreement between the predicted segmentation masks and the corresponding ground-truth annotations [50, 51]. Since the model generates slice-wise 2D predictions, the resulting segmentation masks were subsequently reconstructed into a 3D volume for final evaluation. All quantitative metrics were then computed on these reconstructed 3D segmentation volumes to ensure a consistent volumetric assessment of segmentation performance.

#### 2.2.1 3D Dice Similarity Coefficient (DSC)

The 3D Dice Similarity Coefficient (DSC) was used as the primary overlap-based metric to evaluate the agreement between the predicted and ground-truth segmentation volumes. It measures the spatial overlap between the two 3D volumes, Equation 27:

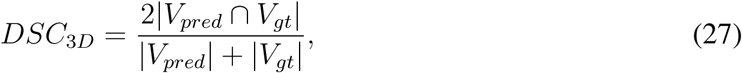

where *V_pred_* and *V_gt_* denote the predicted and ground-truth segmentation volumes, respectively. The term |*V_pred_* ∩ *V_gt_*| represents the number of overlapping voxels (3D pixels) between the two volumes, while |*V_pred_*| and |*V_gt_*| represent the total number of voxels in the predicted and ground-truth segmentations. The 3D DSC ranges from 0 to 1, where a value of 1 indicates perfect volumetric overlap and a value of 0 indicates no overlap between the predicted and ground-truth segmentations.

#### 2.2.2 Intersection over Union (IoU)

The Intersection over Union (IoU), also known as the Jaccard Index, was used to quantify the overlap between the predicted and ground-truth segmentation volumes. It measures the ratio of the intersection to the union of the two 3D volumes, as shown in Equation 28:

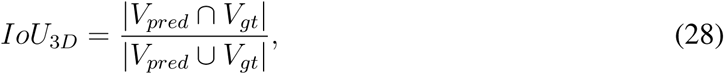

where the term |*V_pred_* ∩ *V_gt_*| represents the number of overlapping voxels, while |*V_pred_* ∪ *V_gt_*| represents the total number of voxels covered by either volume. The IoU ranges from 0 to 1, where a value of 1 indicates perfect volumetric overlap and a value of 0 indicates no overlap between the predicted and ground-truth segmentations.

#### 2.2.3 Sensitivity (Recall)

Sensitivity (Recall) measures the proportion of true positive voxels that are correctly identified by the segmentation model in the 3D volume. It evaluates how well the model detects the ground-truth structure as shown in Equation 29:

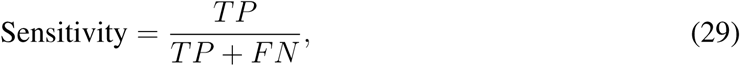

where *TP* and *FN* denote the number of true positive and false negative voxels in the 3D segmentation volume, respectively. Sensitivity ranges from 0 to 1, where a value of 1 indicates that all ground-truth voxels are correctly identified (no false negatives), while a value of 0 indicates that none of the true positive voxels are detected.

#### 2.2.4 95th Percentile Hausdorff Distance (HD95)

To reduce the influence of outlier boundary points, the 95th percentile Hausdorff Distance (HD95) is used as a more robust evaluation metric for 3D segmentation performance. It measures the boundary discrepancy between the predicted segmentation volume and the ground truth volume. HD95 is computed using the 95th percentile of all shortest voxel-to-surface distances in both directions, i.e., from the predicted volume to the ground truth and from the ground truth to the predicted volume. The final HD95 value is obtained by taking the maximum of these two percentile distances, Equation 30:

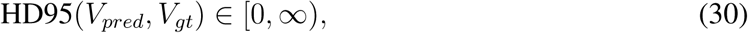

where a value of 0 indicates perfect boundary agreement, while larger values indicate greater segmentation error.

#### 2.2.5 Relative Volume Error (RVE)

The Relative Volume Error (RVE) is used to evaluate the accuracy of volumetric estimation. This metric is a measurement of the relative difference between the predicted and ground truth volumes and is shown in Equation 31:

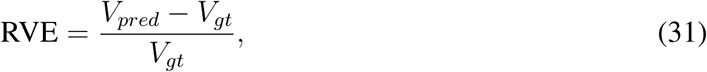

where *V_pred_* and *V_gt_* denote the predicted and ground truth volumes, respectively. Values of RVE close to zero indicate accurate volume estimation, while positive or negative values indicate overestimation or underestimation of the true volume.

## 3 Results and Discussion

In this study, three different backbone networks were evaluated for the proposed DeepLabV3-based segmentation framework, namely ResNet-50, ResNet-101, and Inception-ResNet-v2. Each backbone was integrated into the DeepLabV3 architecture to assess its impact on segmentation performance and feature extraction. Both 2D and 2.5D CNN architectures were investigated to evaluate the effect of incorporating spatial contextual information on segmentation accuracy. For the 2D models, single slices were used as input, whereas for the 2.5D models, a stack of adjacent slices was employed to include inter-slice contextual information. The proposed framework was trained and evaluated using a dataset consisting of 1000 DICOM slices per lizard claw, acquired across three anatomical planes, namely axial, coronal, and sagittal views. To improve model generalization and reduce overfitting, data augmentation was applied, increasing the effective dataset size from 1000 to 5000 slices. For ResNet-50 and ResNet-101, the input image size was set to 224 × 224, while for Inception-ResNet-v2, an input size of 299 × 299 was used to match its native architectural requirements.

To ensure a reliable evaluation under limited data conditions, the dataset consisting of 14 claws was partitioned at the subject level. In each experimental run, 9 claws were used for training, 2 claws were used for validation, and 2 claws were reserved as an internal test set for model assessment during development. Additionally, a leave one out cross validation (LOOCV) strategy was employed, where one claw at a time was held out as an external test set to evaluate the generalisation performance of the model [52]. All DICOM slices belonging to a given claw were strictly confined to their respective split to avoid data leakage. This subject level partitioning ensures that the reported performance reflects true anatomical generalisation across unseen claws.

All experiments were implemented and trained using MATLAB R2025b. Training was performed on a system equipped with a 16-core CPU and 16 GB of RAM. The same hardware configuration was used consistently across all experiments to ensure a fair comparison between the different backbone networks.

### 3.1 Superpixel Based Annotation

The proposed superpixel-based segmentation was applied to CT slices with a resolution of 256×256 pixels. To preserve fine anatomical details and ensure accurate delineation of tissue boundaries, each slice was initially over-segmented into 2000 superpixels. During parameter tuning, the compactness parameter was initially set to 5 and subsequently reduced to 1 based on visual assessment of the segmentation quality. This approach helped ensure that the optimal segmentation integer was used in our work. The lower compactness value allowed the generated superpixels to better conform to the irregular and highly curved boundaries observed within the lizard claw structures, thereby improving the representation of bone and keratin regions.

Following the over-segmentation stage, adjacent superpixels were merged according to the valley points identified in the intensity histograms of the segmented regions. This merging strategy grouped superpixels with similar intensity characteristics into larger homogeneous regions while preserving the underlying anatomical structures. The resulting regions were then classified into background, keratin, and bone classes using literature-based intensity thresholds. Specifically, intensity values between 0 and 20 were assigned to the background class, values between 20 and 200 were classified as keratin, and values greater than 200 were classified as bone.

The segmentation results obtained using the proposed superpixel merging and threshold-based classification approach are presented in Figure 4. As shown in the figure, the method successfully delineates the bone and keratin components of the lizard claw while maintaining anatomically consistent boundaries. The generated segmentations were subsequently used as the ground-truth annotations for training and evaluating the deep learning models. In the absence of manually annotated ground-truth segmentations for the lizard claw, the validity of the proposed superpixel-based segmentation approach was evaluated using literature-derived intensity thresholds for bone and keratin classification [42–45, 53], together with qualitative visual inspection of the segmented anatomical structures.

**Figure 4:**
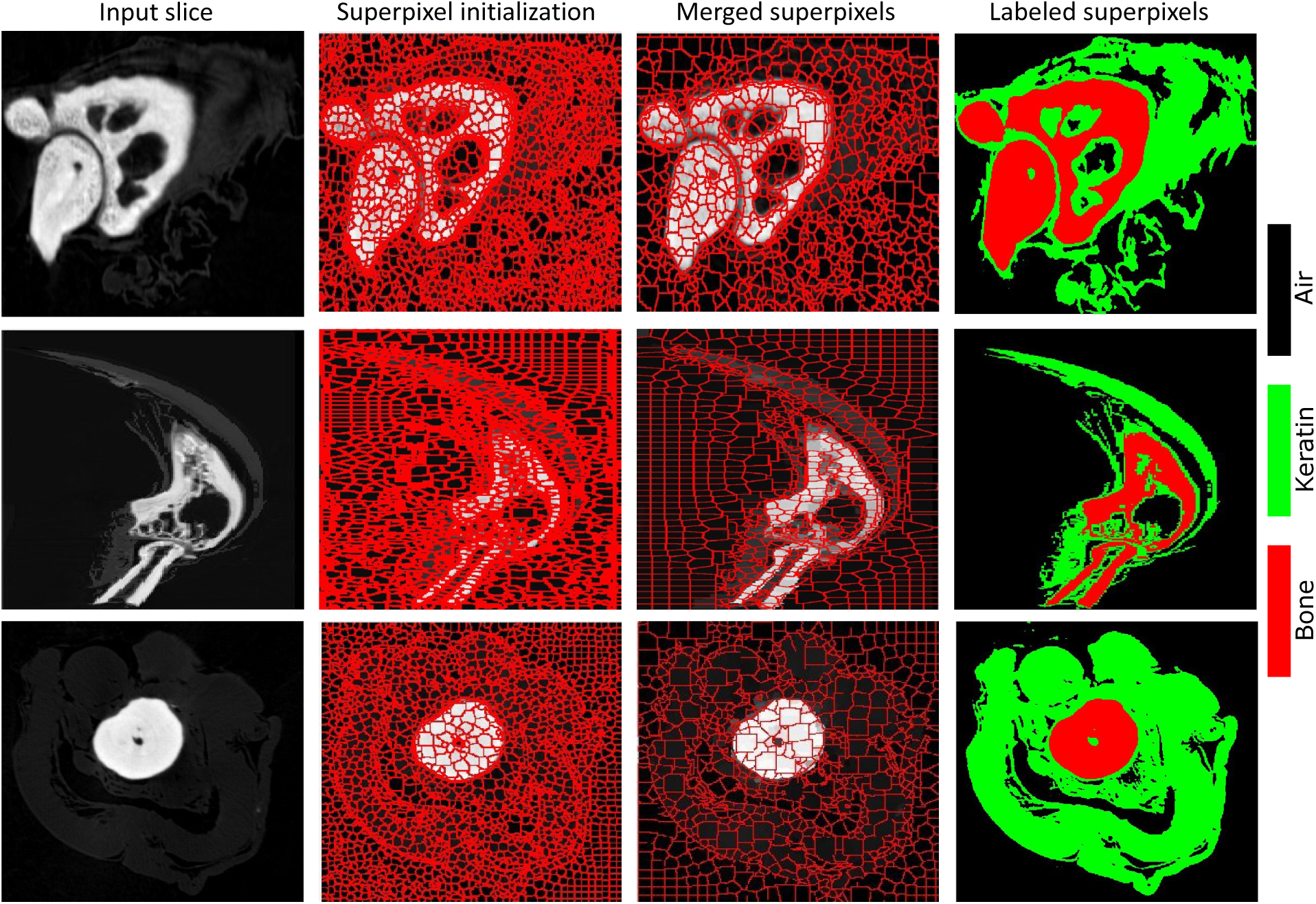
Illustration of the superpixel based segmentation of the lizard claw CT slice showing the delineation of bone and keratin regions. The segmentation was performed using superpixel merging based on histogram intensity distributions, followed by classification using literature-based intensity thresholds for bone and keratin [42–45, 53]. This approach groups homogeneous regions and assigns tissue labels according to their intensity characteristics, resulting in accurate anatomical boundary delineation. The final segmentation is used as the ground-truth annotation for training and evaluation of the deep learning models

After performing the proposed superpixel-based annotation approach, the percentage composition of bone and keratin tissues was quantified for each lizard claw specimen. As shown in Figure 5, the resulting analysis presents the relative distribution of mineralized (bone) and keratinized tissues across the dataset. This quantitative representation highlights the variability in tissue composition among individual claws, reflecting inherent anatomical differences. Importantly, this composition distribution is used as the ground-truth reference for evaluating and predicting tissue composition percentages generated by the deep learning models.

**Figure 5:**
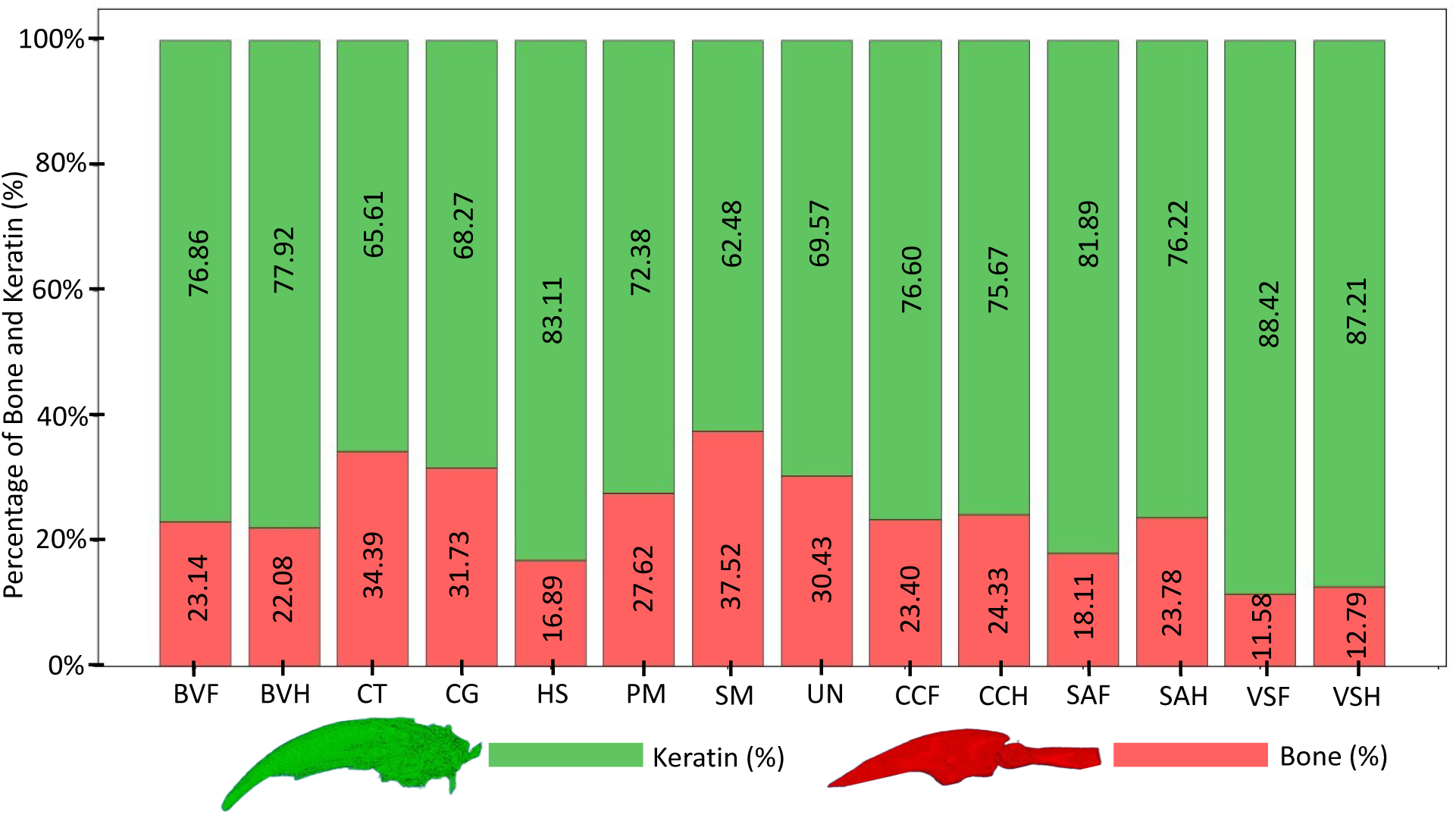
Figure illustrates the percentage composition of bone and keratin within each lizard claw specimen, as determined by the proposed superpixel-based segmentation method. The segmented bone and keratin regions were subsequently used to generate the ground-truth annotations for training and evaluation of the deep learning models. The graph highlights the relative distribution of mineralized (bone) and keratinized tissues across the dataset and demonstrates the variability in tissue composition among individual claws

#### 3.1.1 Performance Analysis of CNN Models

Figure 6 presents a comprehensive comparison of validation accuracy for three backbone networks ResNet-50, ResNet-101, and Inception-ResNet-v2 evaluated under both 2D and 2.5D CNN frameworks across different combinations of learning rates and batch sizes. Overall, the results demonstrate a consistent improvement in performance when moving from 2D to 2.5D architectures, confirming the benefit of incorporating inter-slice contextual information for lizard claw CT segmentation. Across all models, the 2.5D CNN variants outperform their 2D counterparts, particularly at optimized hyperparameter settings.The corresponding training and validation learning curves, including accuracy and loss plots for all evaluated models, are provided in the Electronic Supplementary Material (ESM) as Figures A.1–A.6.

**Figure 6:**
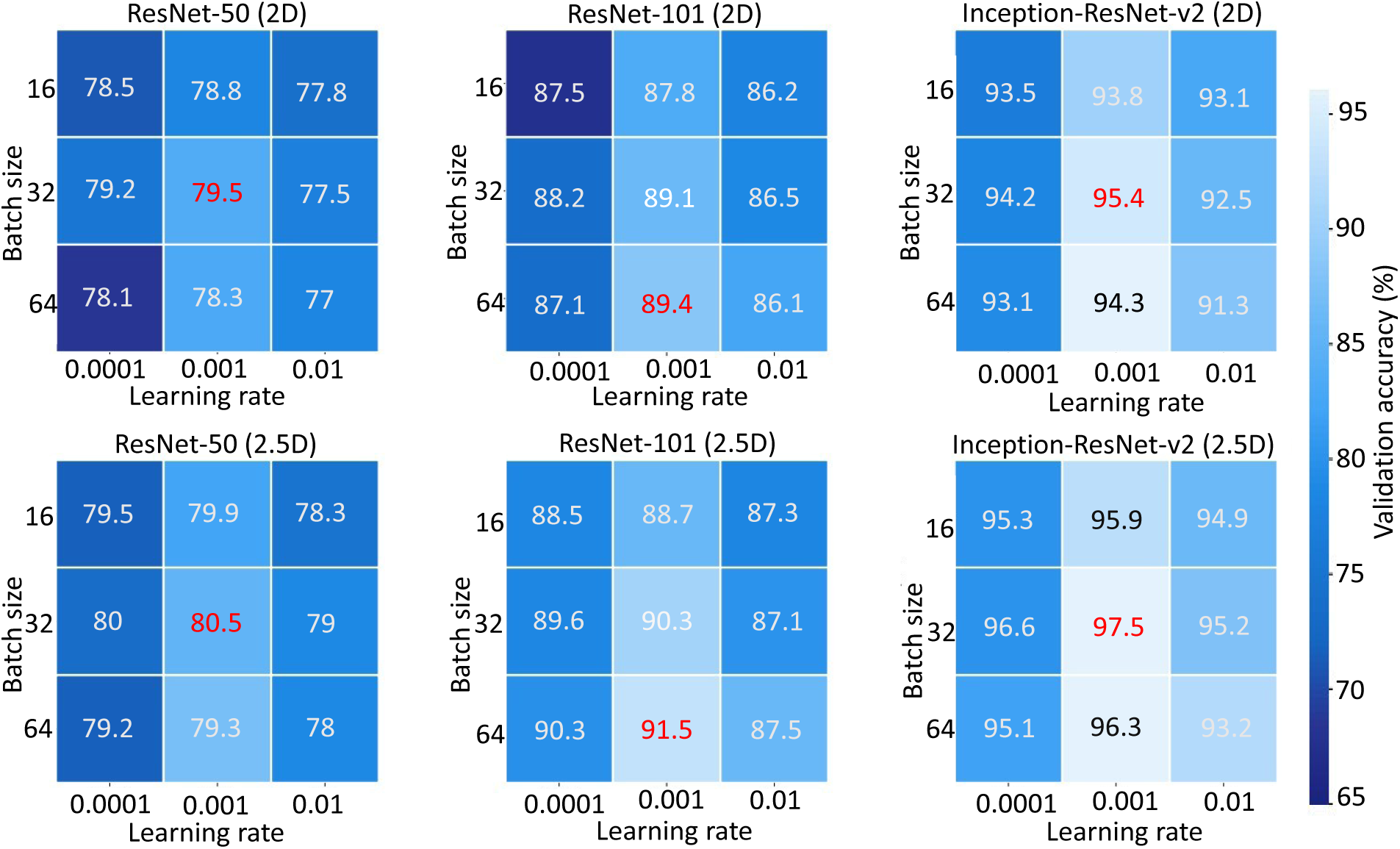
Illustration of validation accuracy (%) comparison of ResNet-50, ResNet-101, and Inception-ResNet-v2 backbones under 2D and 2.5D CNN frameworks across different learning rates (0.0001, 0.001, 0.01) and batch sizes (16, 32, 64). The heatmaps illustrate the effect of hyperparameter selection on model performance. The results demonstrate that 2.5D models consistently outperform 2D models, with Inception-ResNet-v2 achieving the highest overall accuracy

Among the evaluated architectures, Inception-ResNet-v2 achieves the highest performance, reaching a peak validation accuracy of 97.5% (2.5D) at a learning rate of 0.001 and batch size of 32, indicating its superior feature extraction capability due to the combination of inception modules and residual connections. In contrast, ResNet-50 demonstartes a comparatively lower performance, with maximum accuracies around 80–81%, reflecting its limited representational depth for this complex segmentation task. ResNet-101 provides a balanced performance, achieving intermediate accuracy between ResNet-50 and Inception-ResNet-v2. Its best performance is observed in the 2.5D configuration, reaching approximately 91.5% accuracy, highlighting the advantage of deeper residual learning for capturing fine anatomical structures. From the hyperparameter perspective, a learning rate of 0.001 consistently yields the best performance across all models, while extremely low (0.0001) or high (0.01) learning rates result in reduced accuracy. Similarly, a batch size of 32 provides the most stable and optimal training behavior, suggesting a good balance between gradient stability and generalization.

Overall, the results confirm that both architectural choice and input dimensionality significantly influence segmentation performance, with Inception-ResNet-v2 (2.5D) providing the most robust and accurate results for lizard claw keratin and bone classification. The superior performance of the Inception-ResNet 2.5D model over the 2D approach can be attributed to its ability to process local volumetric data, which provides a stronger and more informative representation of anatomical structures. Unlike 2D models that analyze each slice independently, the 2.5D architecture takes multiple adjacent slices as input channels, allowing it to capture inter-slice dependencies and preserve spatial continuity across the volume. This enables the model to learn richer contextual features that better reflect the true 3D structure of the anatomy. As a result, the incorporation of local volumetric information leads to improved feature representation and demonstrates that the 2.5D approach performs better than the 2D model, particularly in medical imaging tasks where relevant patterns are not confined to a single slice.

Figures 7, 8, and 9 illustrate the segmentation performance of different CNN architectures on axial, coronal, and sagittal CT slices, respectively, of a lizard claw *Basiliscus vittatus* (hind limb). The input stack, ground truth, and predicted masks are compared across ResNet-50, ResNet-101, and Inception-ResNet-v2 models implemented in both 2D and 2.5D frameworks. Keratin and bone regions are effectively highlighted in the predictions, demonstrating each model’s capability to preserve anatomical structure and boundary continuity. The results demonstrate that 2.5D Inception-ResNet-v2 shows the most accurate and stable segmentation, closely matching the ground truth annotations as comapre to other architectures.

**Figure 7:**
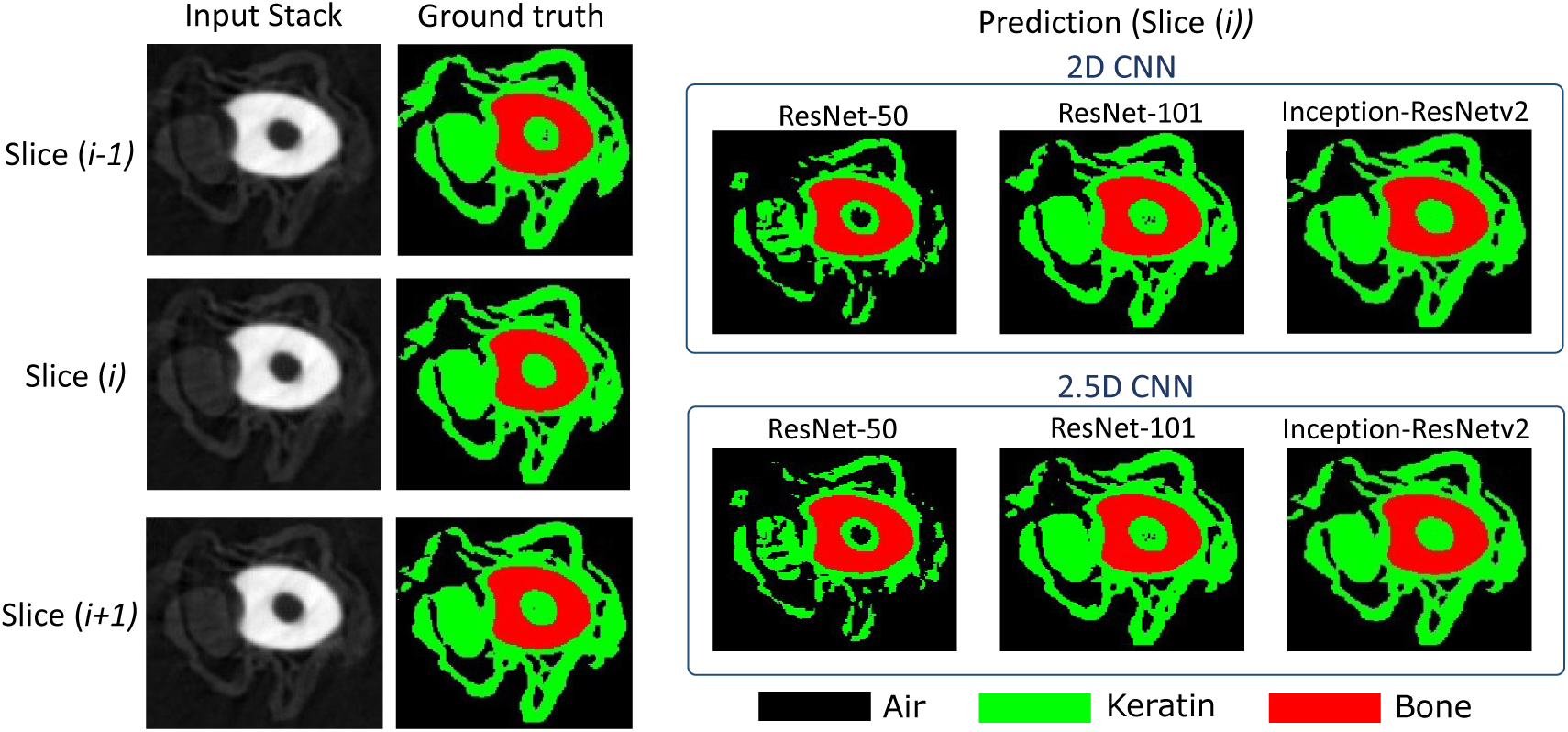
Axial CT slice of a lizard claw (*Basiliscus vittatus* (hind)) showing keratin and bone segmentation results. The ground-truth annotation is compared with predictions obtained using ResNet-50, ResNet-101, and Inception-ResNet-v2 architectures implemented in both 2D and 2.5D CNN frameworks. For the 2.5D models, adjacent slices are used as contextual input, with the target segmentation corresponding to slice(i+1). Keratin and bone regions highlight each model’s ability to delineate tissue boundaries and preserve anatomical structures

**Figure 8:**
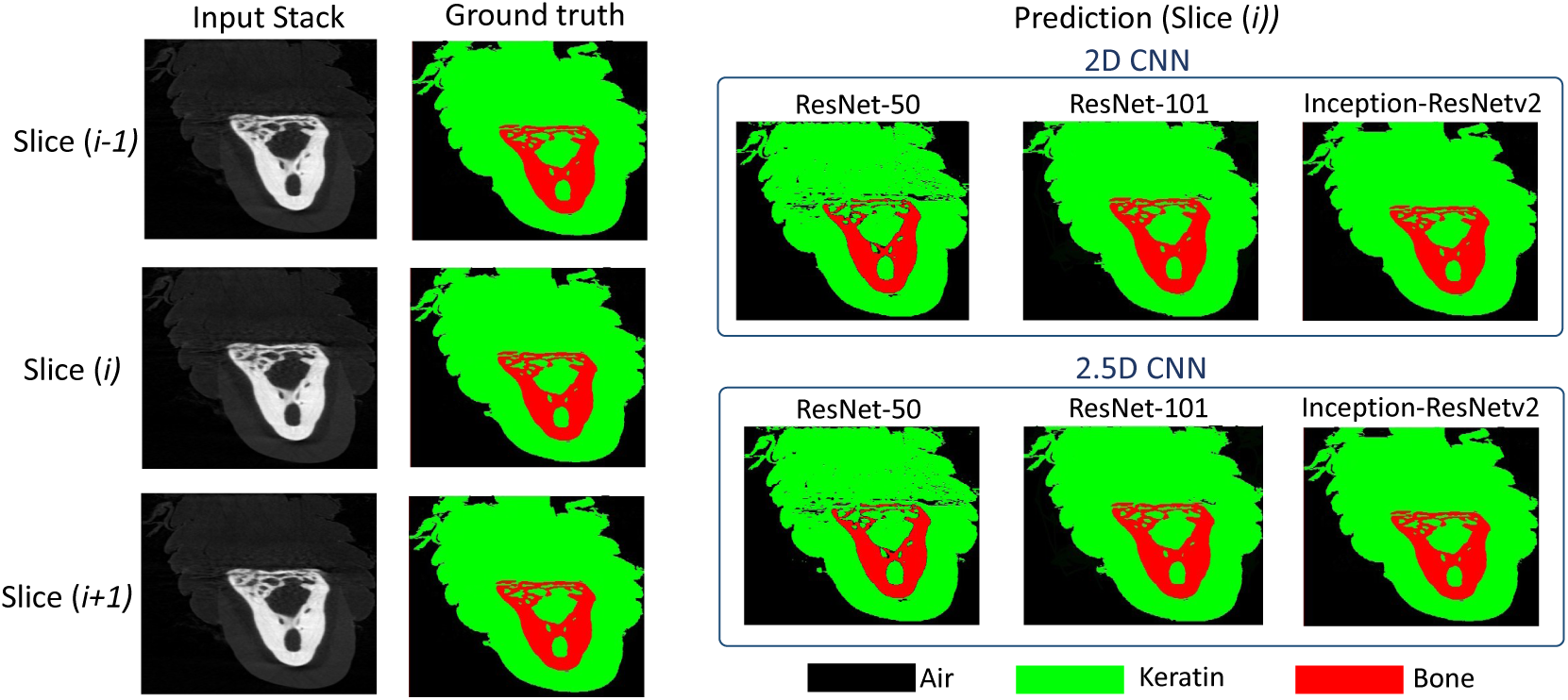
Coronal CT slice of a lizard claw (*Basiliscus vittatus* (hind)) showing keratin and bone segmentation results. The ground-truth annotation is compared with predictions obtained using ResNet-50, ResNet-101, and Inception-ResNet-v2 architectures implemented in both 2D and 2.5D CNN frameworks. For the 2.5D models, adjacent slices are used as contextual input, with the target segmentation corresponding to slice(i+1). Keratin and bone regions highlight each model’s ability to delineate tissue boundaries and preserve anatomical structures

**Figure 9:**
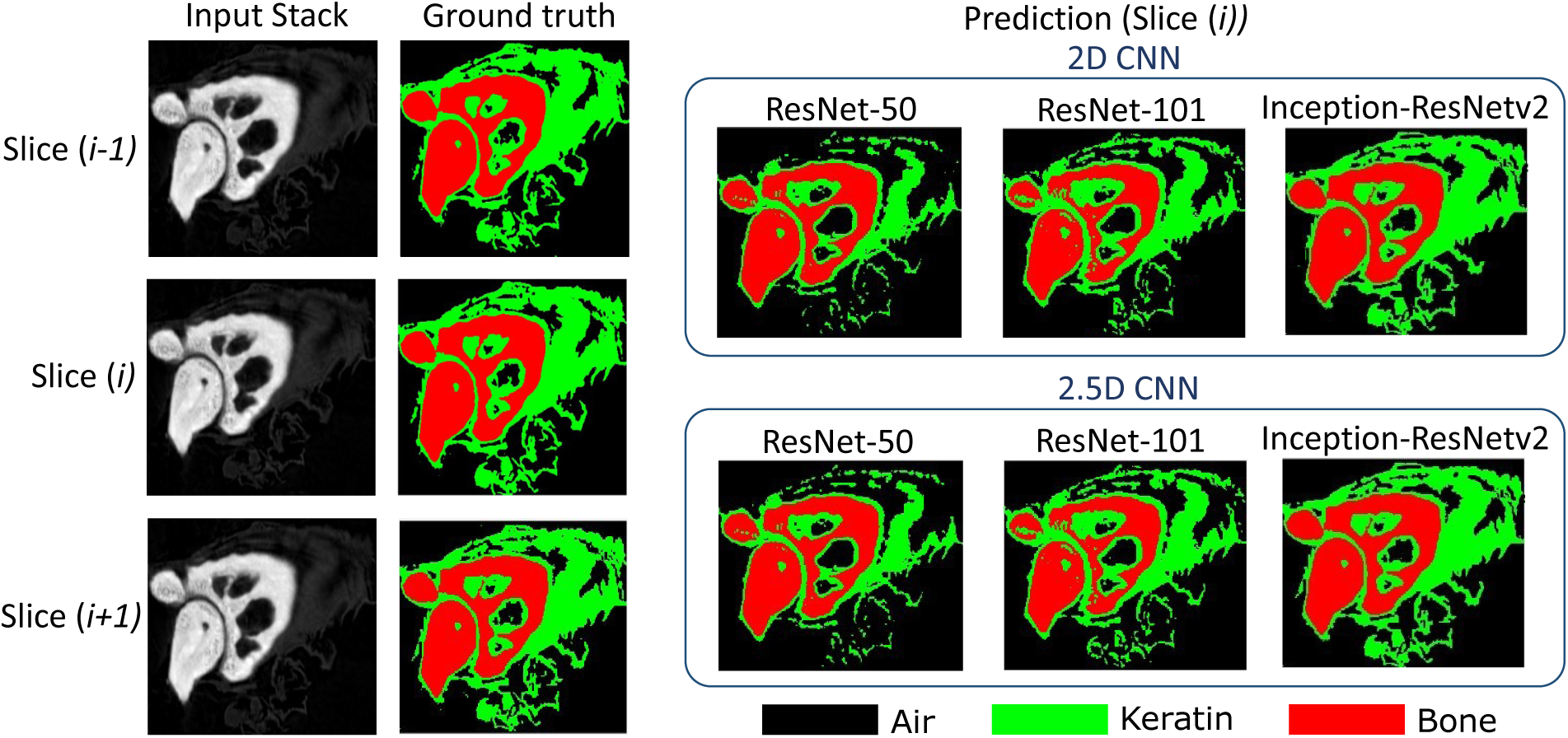
Sagittal CT slice of a lizard claw (*Basiliscus vittatus* (hind)) showing keratin and bone segmentation results. The ground-truth annotation is compared with predictions obtained using ResNet-50, ResNet-101, and Inception-ResNet-v2 architectures implemented in both 2D and 2.5D CNN frameworks. For the 2.5D models, adjacent slices are used as contextual input, with the target segmentation corresponding to slice (i+1). Keratin and bone regions highlight each model’s ability to delineate tissue boundaries and preserve anatomical structures

#### 3.1.2 Performance Assessment using Evaluation Metrics

Table 2 provides information on the performance of the evaluated 2D and 2.5D CNN architectures for 3D segmentation after reconstructing slice-wise predictions into volumetric segmentations. Among the 2D architectures, Inception-ResNet-v2 achieved the highest performance, with a DSC of 0.94 ± 0.03, IoU of 0.92 ± 0.04, and Sensitivity of 0.90 ± 004, while also producing the lowest HD95 0.31 ± 0.03 and RVE 0.34 ± 0.04. ResNet101 demonstrated moderate performance (DSC = 0.86 ± 0.04, IoU = 0.75 ± 0.05), whereas ResNet50 yielded the lowest segmentation accuracy, with a DSC of 0.74 ± 0.06 and IoU of 0.63 ± 0.07. These findings suggest that deeper and more advanced feature extraction architectures provide superior representation of anatomical structures, leading to improved segmentation quality. A similar trend was observed for the 2.5D models. The 2.5D Inception-ResNet-v2 achieved the best overall performance, obtaining the highest DSC (0.96 ± 0.02), IoU (0.95 ± 0.03), and Sensitivity (0.93 ± 0.03), together with the lowest HD95 (0.28 ± 0.02) and RVE (0.30 ± 0.02). Compared with its 2D counterpart, the model showed improvements of approximately 2 percentage points in DSC, 3 percentage points in IoU, and 3 percentage points in Sensitivity, while reducing HD95 and RVE. Likewise, the 2.5D ResNet101 improved upon the 2D version, increasing DSC from 0.86 to 0.89 and reducing RVE from 0.49 to 0.38. Although the performance gain for ResNet50 was smaller, consistent improvements were still observed across all metrics.

**Table 2:** Performance comparison of 2D and 2.5D models for 3D segmentation. Slice-wise predictions generated by the 2D and 2.5D CNN architectures were reconstructed into a 3D volume and subsequently compared with the corresponding 3D ground-truth segmentation. Evaluation metrics include Dice Similarity Coefficient (DSC), Intersection over Union (IoU), Sensitivity, 95th-percentile Hausdorff Distance (HD95), and Relative Volume Error (RVE). Higher values indicate better performance for DSC, IoU, and Sensitivity, whereas lower values indicate better performance for HD95 and RVE.

| Model | DSC | IoU | Sensitivity | HD95 | RVE |
| --- | --- | --- | --- | --- | --- |
| **2D Models** |  |  |  |  |  |
| 2D ResNet50 | $0.72 \pm 0.06$ | $0.63 \pm 0.07$ | $0.75 \pm 0.06$ | $0.62 \pm 0.05$ | $0.65 \pm 0.07$ |
| 2D ResNet101 | $0.86 \pm 0.04$ | $0.75 \pm 0.05$ | $0.87 \pm 0.05$ | $0.46 \pm 0.04$ | $0.49 \pm 0.04$ |
| 2D Inception-ResNet-v2 | $0.94 \pm 0.03$ | $0.92 \pm 0.04$ | $0.90 \pm 0.04$ | $0.31 \pm 0.03$ | $0.34 \pm 0.04$ |
| **2.5D Models** |  |  |  |  |  |
| 2.5D ResNet50 | $0.74 \pm 0.05$ | $0.64 \pm .07$ | $0.77 \pm 0.05$ | $0.60 \pm .05$ | $0.63 \pm 0.06$ |
| 2.5D ResNet101 | $0.89 \pm 0.03$ | $0.78 \pm 0.04$ | $0.90 \pm 0.04$ | $0.42 \pm 0.03$ | $0.38 \pm 0.04$ |
| 2.5D Inception-ResNet-v2 | $0.96 \pm 0.02$ | $0.95 \pm 0.03$ | $0.93 \pm 0.03$ | $0.28 \pm .02$ | $0.30 \pm 0.02$ |

The lower HD95 values achieved by the 2.5D models indicate more accurate boundary localization and reduced segmentation errors along object contours. Similarly, the reduction in RVE suggests improved estimation of target volume, which is particularly important for quantitative analyses. The superior performance of the 2.5D architectures can be attributed to their ability to exploit contextual information from neighboring slices while maintaining lower computational complexity than fully 3D networks. Overall, the results demonstrate that incorporating inter-slice contextual information through a 2.5D framework enhances segmentation performance. Among all evaluated models, the 2.5D Inception-ResNet-v2 provided the most accurate and robust 3D segmentation, making it the most suitable architecture for the proposed task.

#### 3.1.3 Quantitative Assessment of Keratin and Bone Composition

Following the selection of the 2.5D Inception-ResNet-v2 model as the optimal segmentation architecture, its ability to quantify claw tissue composition was evaluated by comparing the predicted percentages of keratin and bone against the corresponding ground-truth values for each claw under the leave one out cross validation (LOOCV) framework. In each fold, one claw was excluded from training and used exclusively for testing, ensuring that tissue composition estimates were obtained from anatomically unseen specimens. Figure 10 elucidates the percentage composition of keratin and bone for all 14 claws. Overall, the predicted tissue proportions closely matched the ground-truth measurements, demonstrating the model’s ability to accurately quantify volumetric tissue composition in addition to producing accurate segmentations.

**Figure 10:**
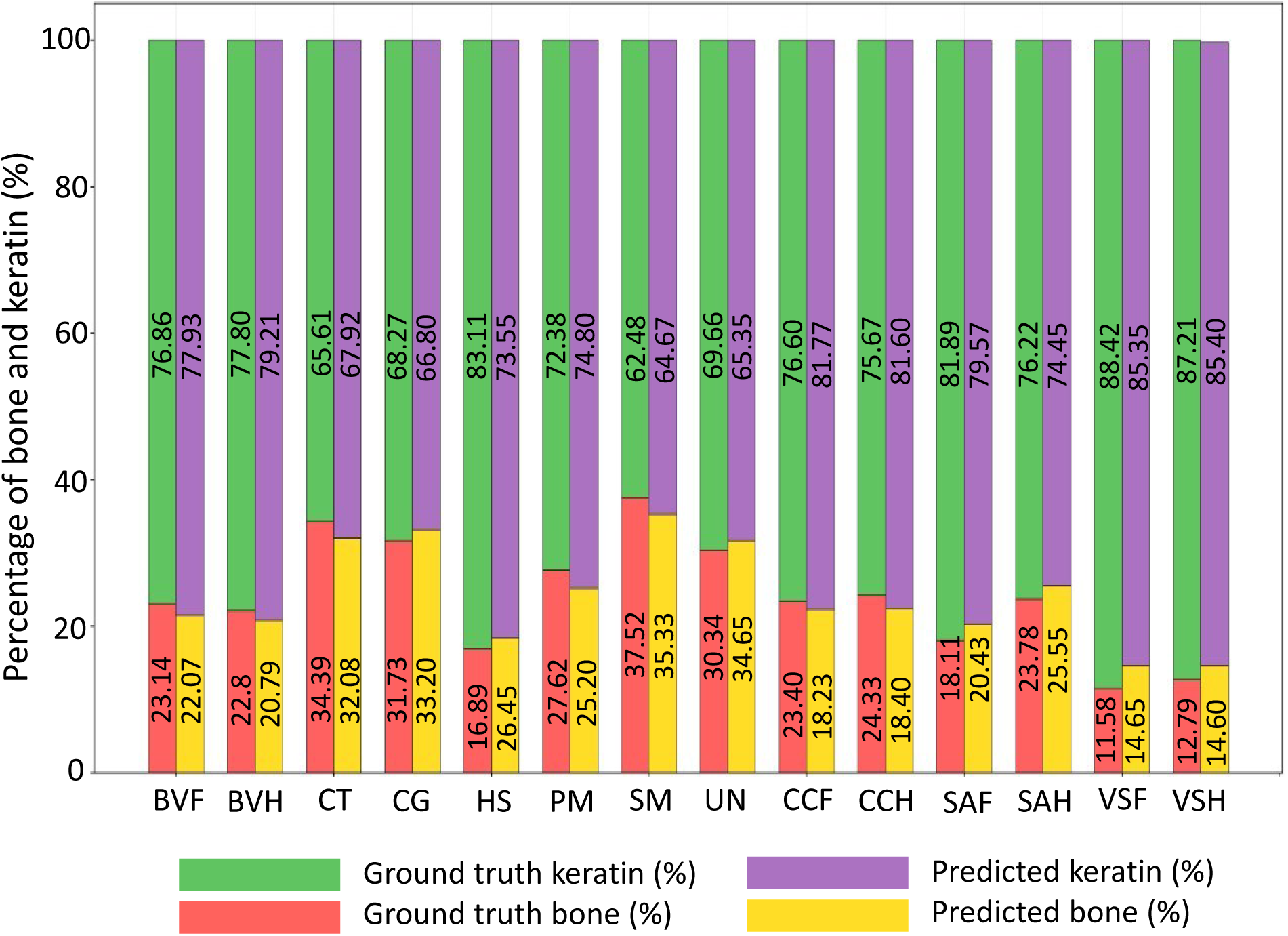
Comparison of ground truth and predicted tissue composition percentages for keratin and bone across all 14 lizard claws under the leave one out cross validation (LOOCV) framework where in each fold, one claw was excluded from training and used exclusively for testing, ensuring that tissue composition estimates were obtained from anatomically unseen specimens. Predictions were generated using the Inception-ResNet-v2 2.5D segmentation model. The close agreement between predicted and ground-truth values demonstrates the model’s ability to accurately quantify volumetric tissue composition

Although minor deviations were observed in some cases, such as slight overestimation or underestimation of bone content in individual claws, these discrepancies remained relatively small and were generally balanced by corresponding changes in keratin percentage. Since keratin and bone together constitute the entire segmented claw volume, small segmentation boundary differences can propagate into percentage calculations, particularly in claws with smaller bone regions. Nevertheless, the close agreement between predicted and ground-truth values across all test claws suggests that the selected model provides reliable quantitative estimates of tissue composition.

These findings demonstrate that the 2.5D Inception-ResNet-v2 model achieved the highest segmentation accuracy among the evaluated architectures and accurately reproduced biologically meaningful measurements of keratin and bone distribution. The strong agreement between predicted and reference tissue percentages supports the suitability of the proposed framework for quantitative anatomical analysis and downstream applications involving volumetric tissue composition assessment. Furthermore, performance may be further improved with access to larger training datasets and a higher number of DICOM slices, which could enhance the model’s ability to generalize and capture more comprehensive spatial context.

The Figures A1–A3 in the Electronic Supplementary Material (ESM) provide a qualitative analysis in which the ground-truth values are compared with the predictions produced by the Inception-ResNet 2.5D model to evaluate the agreement between observed and predicted volumes.

## 4 Conclusions

This study presents a deep learning-based framework for the automated segmentation of bone and keratinous tissue in lizard claws from high-resolution CT imaging data, followed by reconstruction for quantitative analysis. Both 2D and 2.5D convolutional neural network (CNN) architectures were evaluated using DeepLabV3 with ResNet-50, ResNet-101, and Inception-ResNet-v2 backbones. The incorporation of inter-slice contextual information in the 2.5D framework significantly improved segmentation performance compared to the 2D baseline. Among all models, the 2.5D Inception-ResNet-v2 achieved the best overall performance, demonstrating superior accuracy in distinguishing structurally similar tissues and achieving a validation accuracy of 97.5%. These results highlight the effectiveness of the proposed approach for precise tissue delineation in complex claw anatomy. From a modeling perspective, this study provides accurate segmentation of internal claw tissues, enabling detailed characterization of material composition and structural organization in lizard claws. Despite these promising results, the study is limited by the relatively small dataset size. The segmentation accuracy could be further improved by incorporating a larger and more diverse dataset, which would enhance model generalization and robustness across varying anatomical structures.

## Supporting information

Electronic Supplementary Material

