## Supplementary material for "Deep-learning based 3D segmentation of heterogeneous lizard claw tissue from CT data": Electronic Supplementary Material

### Supplementary Material for:
### Deep-learning based 3D segmentation of heterogeneous lizard claw tissue from CT data

Haleema Sadia[1], Kate Manges Douglas,[2] Asyiah Bray,[2]

Andrea Rummel,[2] Parvez Alam,[1*]

[1]School of Engineering, The University of Edinburgh, Edinburgh, UK

[2]Department of BioSciences, Rice University, Houston, TX 77005, USA

July 27, 2026

### A CNN Training Graphs

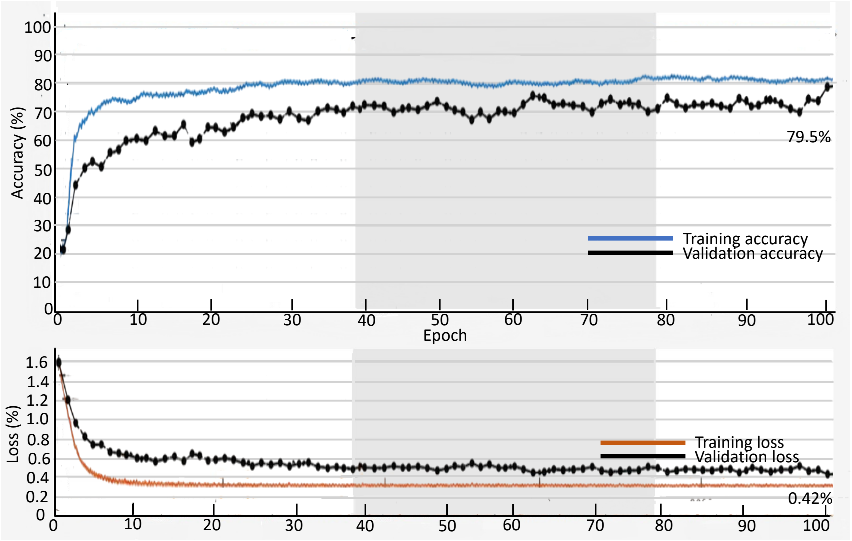


Figure A.1: Training and validation performance of the ResNet-50 2D CNN model over successive epochs, achieving a maximum validation accuracy of 79.5% with a minimum validation loss of 0.42%

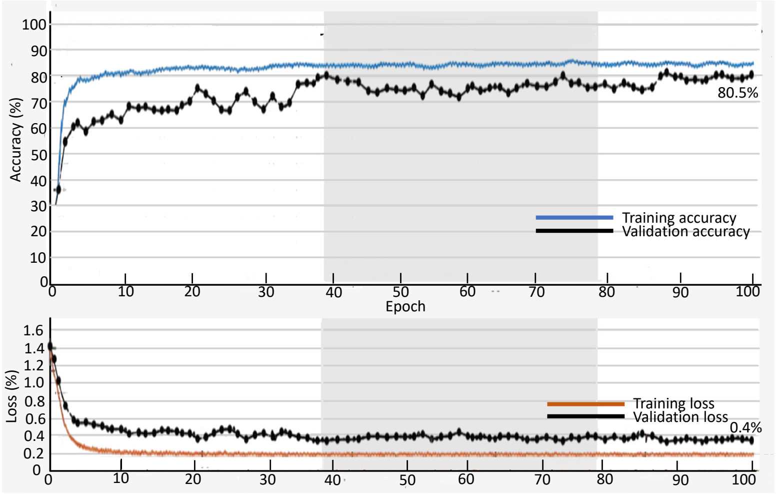


Figure A.2: Training and validation performance of the ResNet-50 2.5D CNN model over successive epochs, achieving a maximum validation accuracy of 80.5% with a minimum validation loss of 0.4%

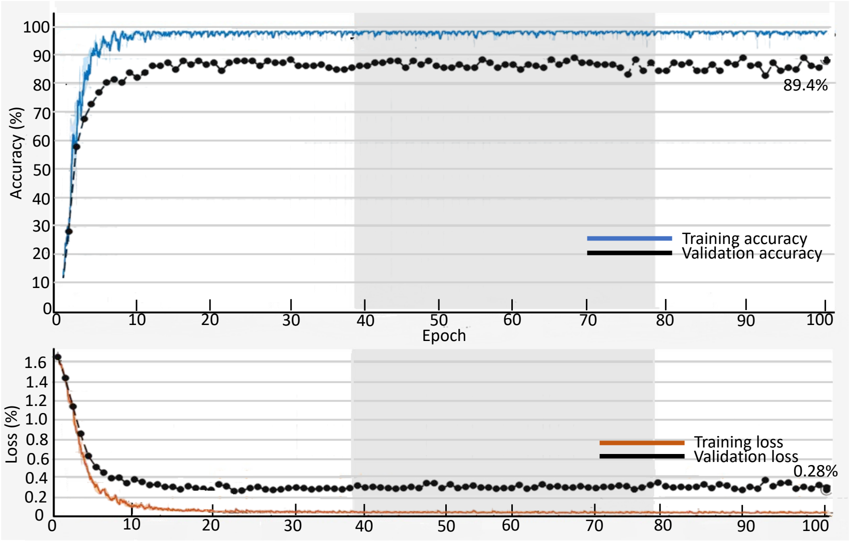


Figure A.3: Training and validation performance of the ResNet-101 2D CNN model over successive epochs, achieving a maximum validation accuracy of 89.4% with a minimum validation loss of 0.28%

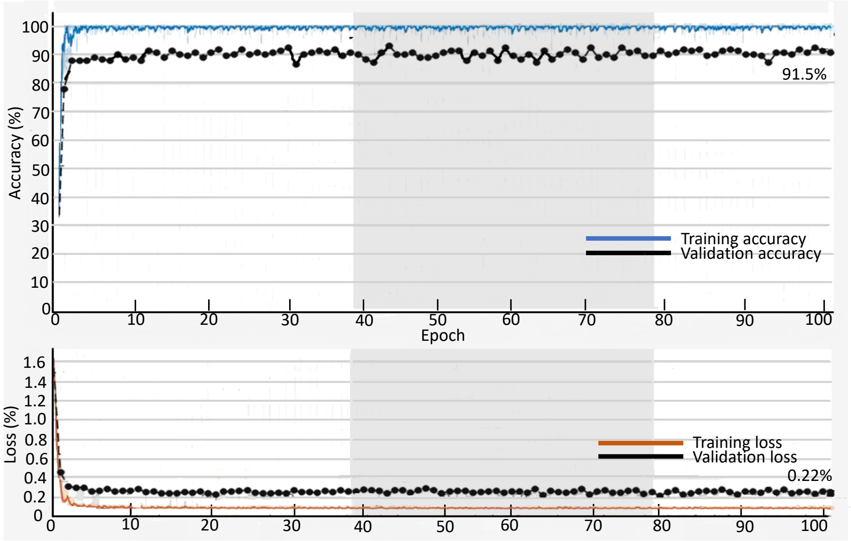


Figure A.4: Training and validation performance of the ResNet-101 2.5D CNN model over successive epochs, achieving a maximum validation accuracy of 91.5% with a minimum validation loss of 0.22%

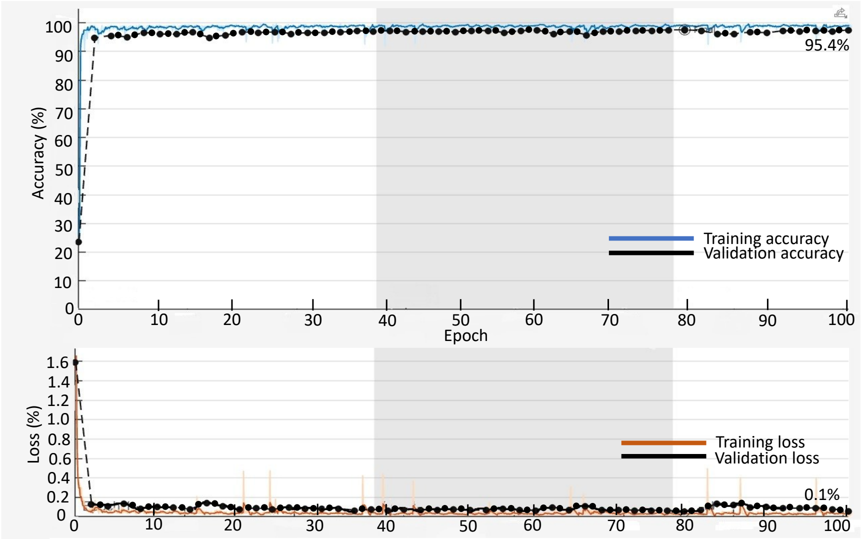


Figure A.5: Training and validation performance of the Inception-ResNet-v2 2D CNN model over successive epochs, achieving a maximum validation accuracy of 95.4% with a minimum validation loss of 0.1%

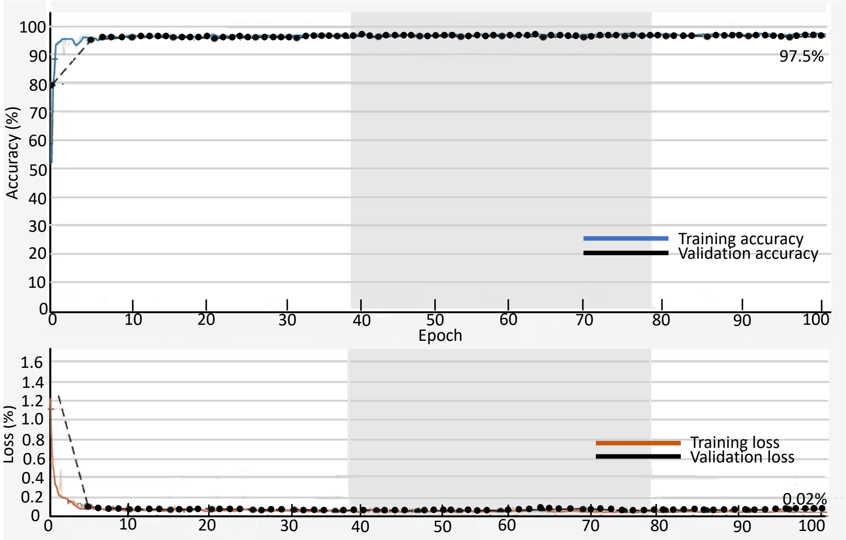


Figure A.6: Training and validation performance of the Inception-ResNet-v2 2.5D CNN model over successive epochs, achieving a maximum validation accuracy of 97.5% with a minimum validation loss of 0.02%

### A Ground Truth vs Inception-ResNet 2.5D Predictions

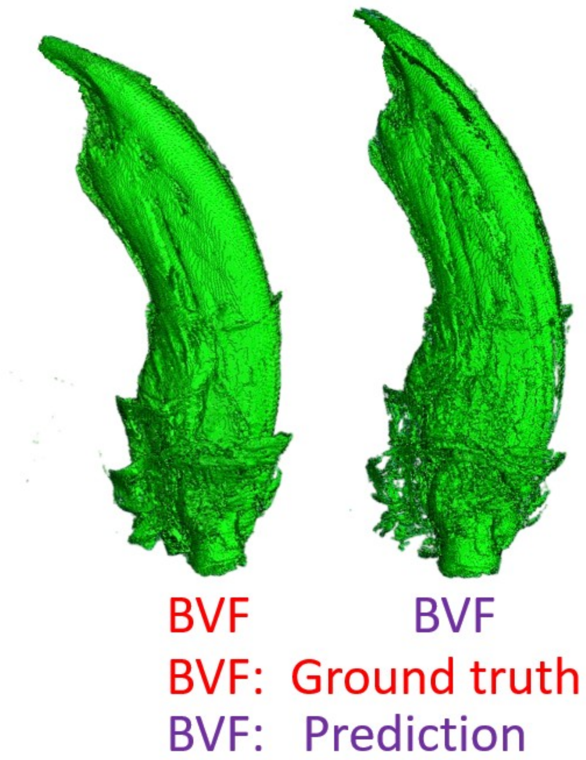


Figure B.1: Illustration of the qualitative comparison between the ground-truth and predicted 3D volume reconstructions of the hind claw of *Basiliscus vittatus* (fore). The predicted volume was generated using the Inception-ResNet 2.5D model under the leave one out cross validation (LOOCV) framework

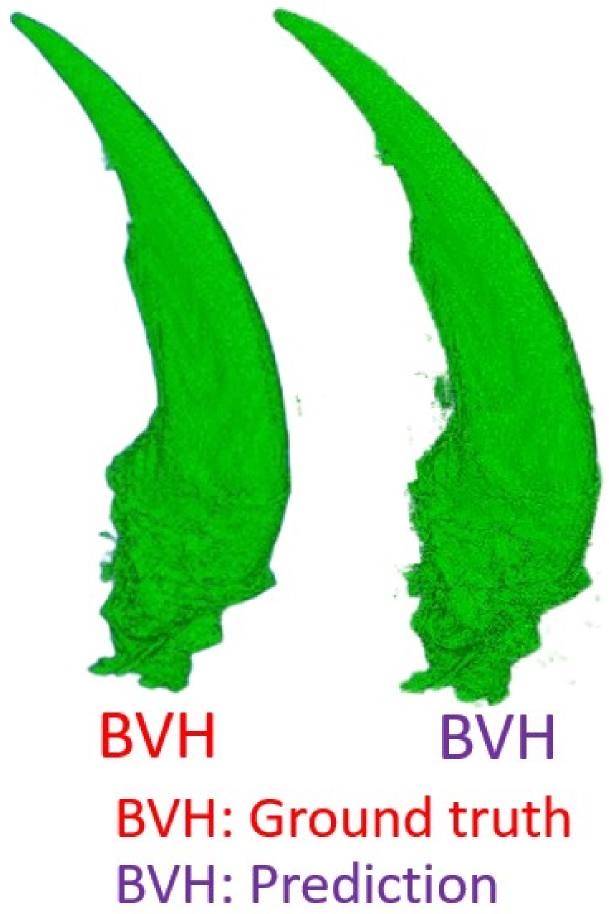


Figure B.2: Illustration of the qualitative comparison between the ground-truth and predicted 3D volume reconstructions of the hind claw of *Basiliscus vittatus* (hind). The predicted volume was generated using the Inception-ResNet 2.5D model under the leave one out cross validation (LOOCV) framework

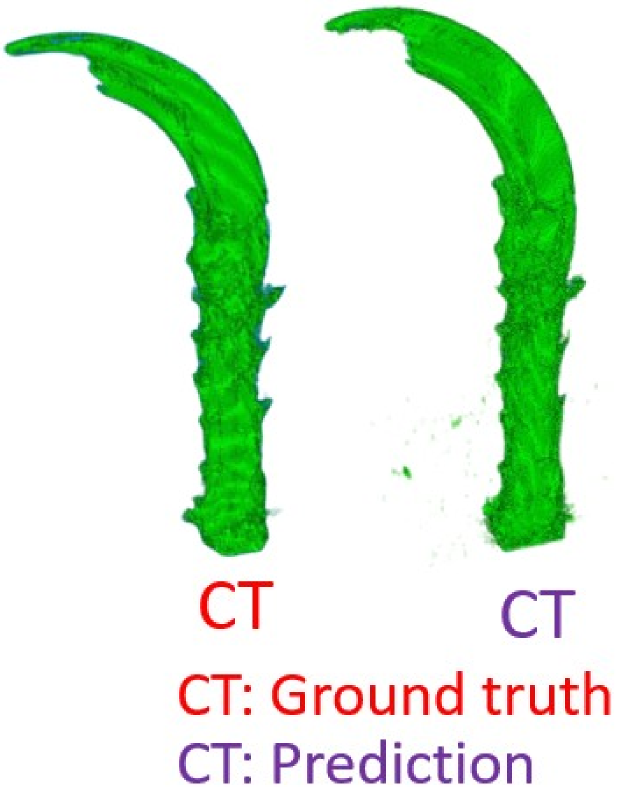


Figure B.3: Illustration of the qualitative comparison between the ground-truth and predicted 3D volume reconstructions of the hind claw of *Cophosaurus texanus*. The predicted volume was generated using the Inception-ResNet 2.5D model under the leave one out cross validation (LOOCV) framework

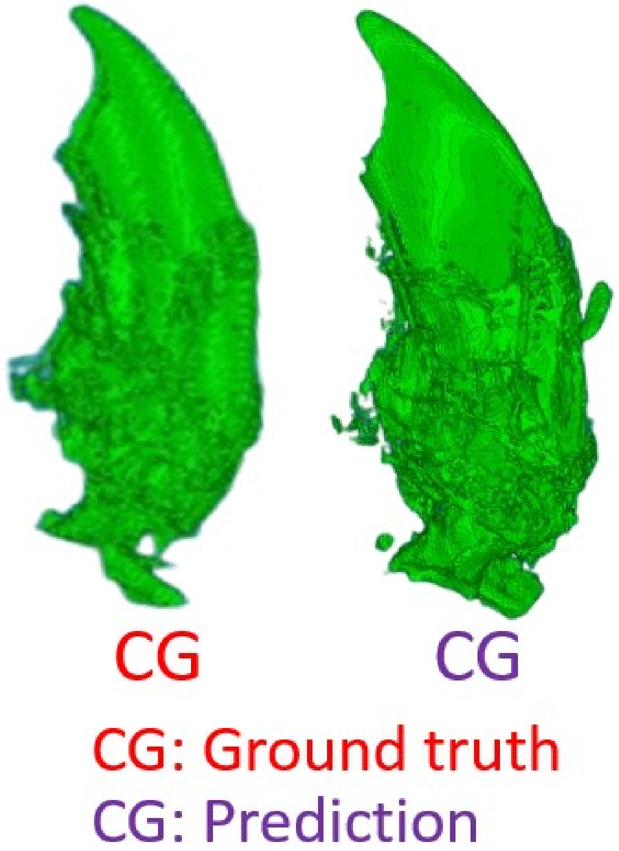


Figure B.4: Illustration of the qualitative comparison between the ground-truth and predicted 3D volume reconstructions of the hind claw of *Cordylus giganteus*. The predicted volume was generated using the Inception-ResNet 2.5D model under the leave one out cross validation (LOOCV) framework

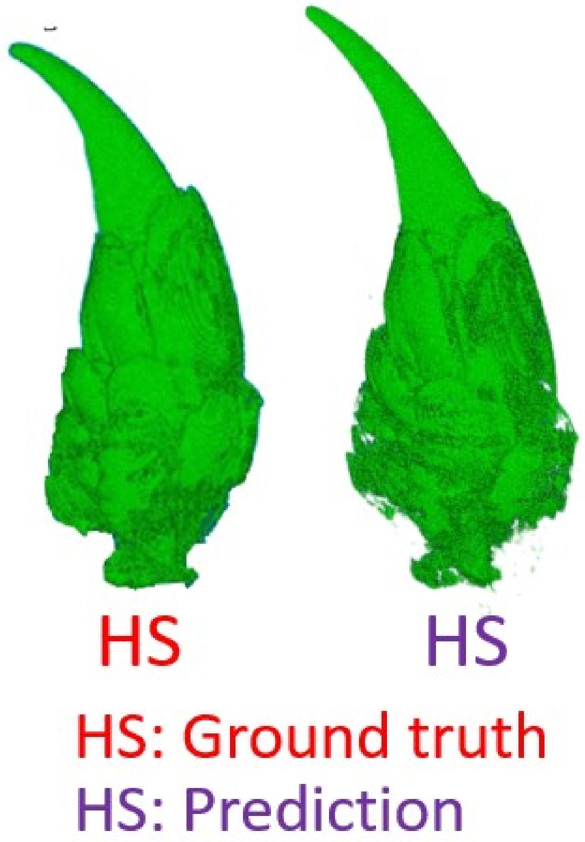


Figure B.5: Illustration of the qualitative comparison between the ground-truth and predicted 3D volume reconstructions of the hind claw of *Heloderma suspectum*. The predicted volume was generated using the Inception-ResNet 2.5D model under the leave one out cross validation (LOOCV) framework

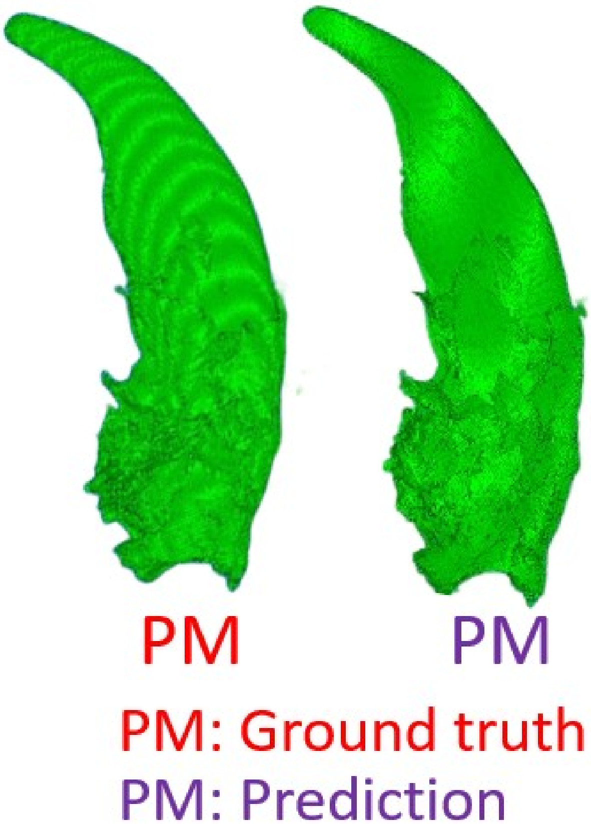


Figure B.6: Illustration of the qualitative comparison between the ground-truth and predicted 3D volume reconstructions of the hind claw of *Phrynosoma modestum*. The predicted volume was generated using the Inception-ResNet 2.5D model under the leave one out cross validation (LOOCV) framework

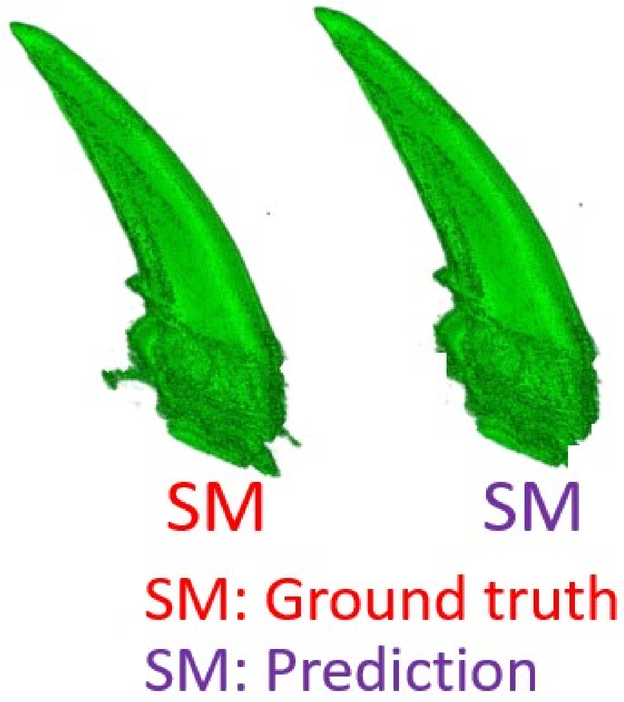


Figure B.7: Illustration of the qualitative comparison between the ground-truth and predicted 3D volume reconstructions of the hind claw of *Salvator merianae*. The predicted volume was generated using the Inception-ResNet 2.5D model under the leave one out cross validation (LOOCV) framework

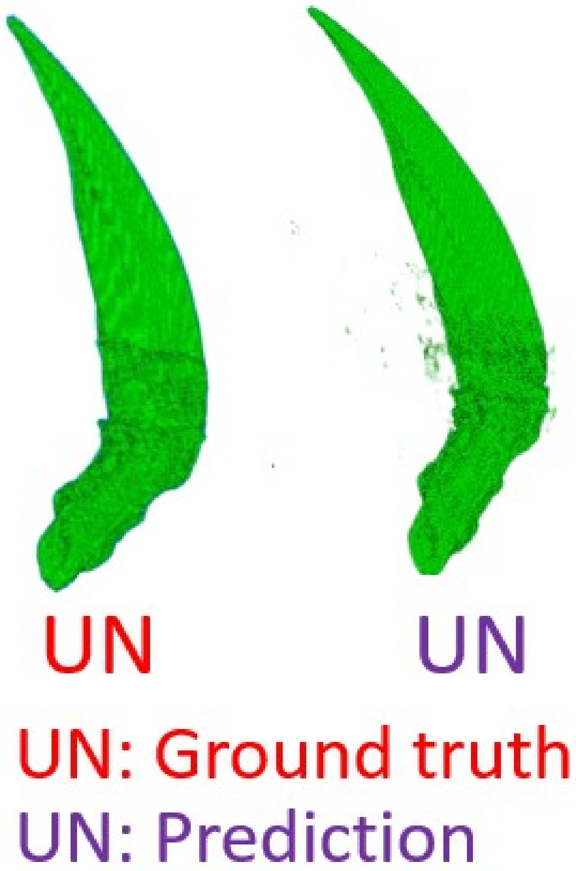


Figure B.8: Illustration of the qualitative comparison between the ground-truth and predicted 3D volume reconstructions of the hind claw of *Uma notata*. The predicted volume was generated using the Inception-ResNet 2.5D model under the leave one out cross validation (LOOCV) framework

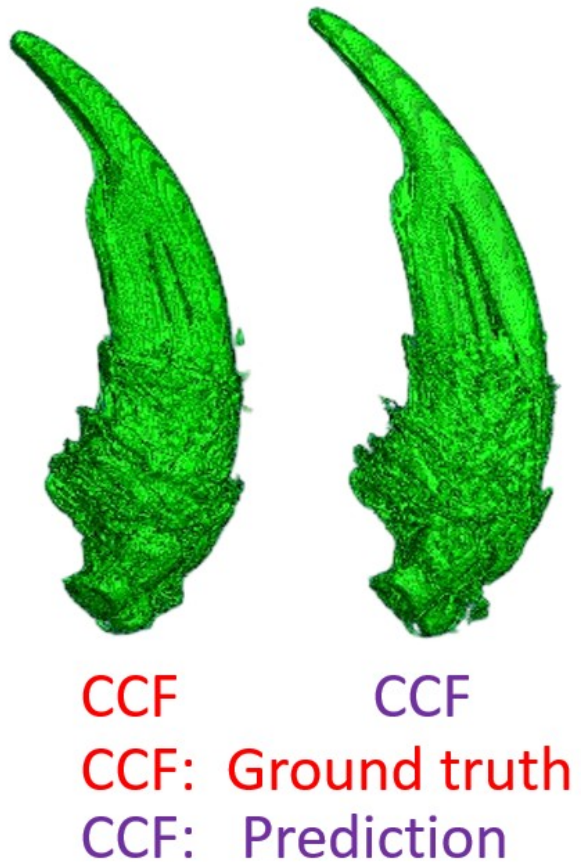


Figure B.9: Illustration of the qualitative comparison between the ground-truth and predicted 3D volume reconstructions of the hind claw of *Crotaphytus collaris* (fore). The predicted volume was generated using the Inception-ResNet 2.5D model under the leave one out cross validation (LOOCV) framework

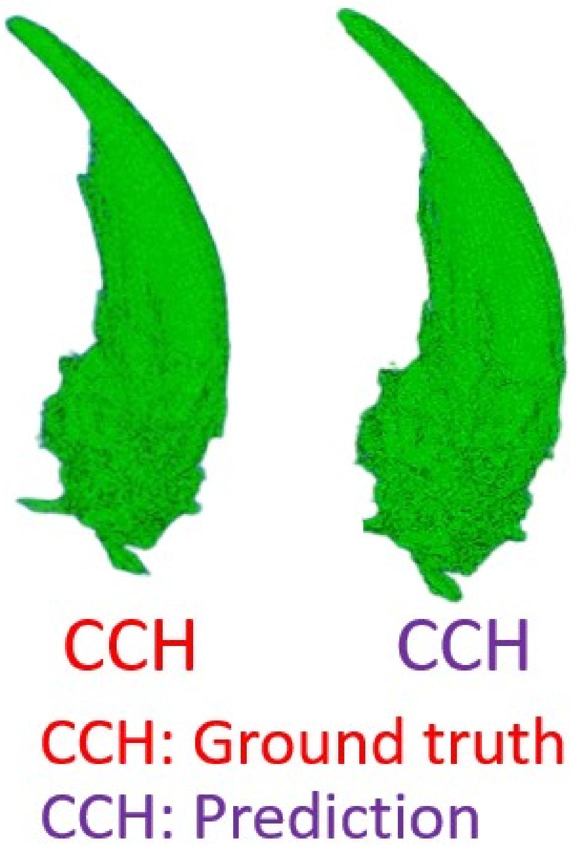


Figure B.10: Illustration of the qualitative comparison between the ground-truth and predicted 3D volume reconstructions of the hind claw of *Crotaphytus collaris* (hind). The predicted volume was generated using the Inception-ResNet 2.5D model under the leave one out cross validation (LOOCV) framework

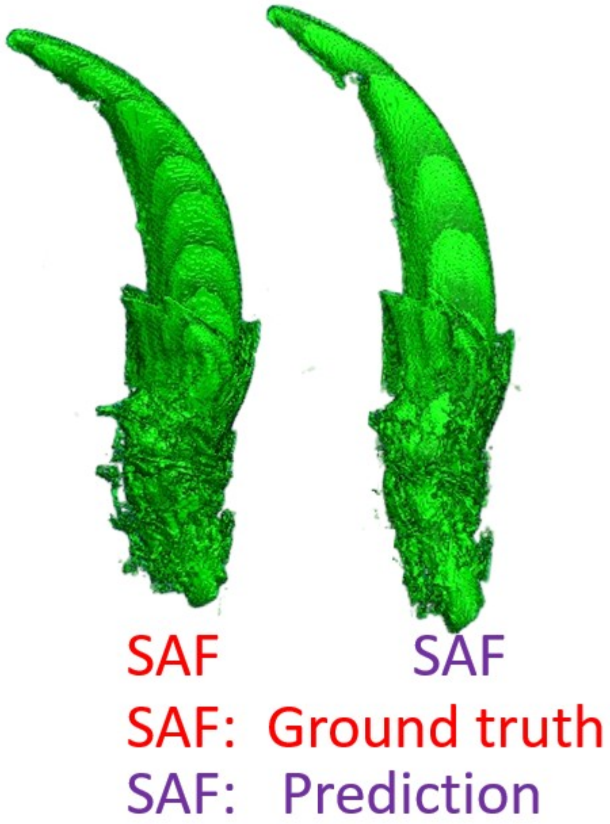


Figure B.11: Illustration of the qualitative comparison between the ground-truth and predicted 3D volume reconstructions of the hind claw of *Sceloporus arenicolus* (fore). The predicted volume was generated using the Inception-ResNet 2.5D model under the leave one out cross validation (LOOCV) framework

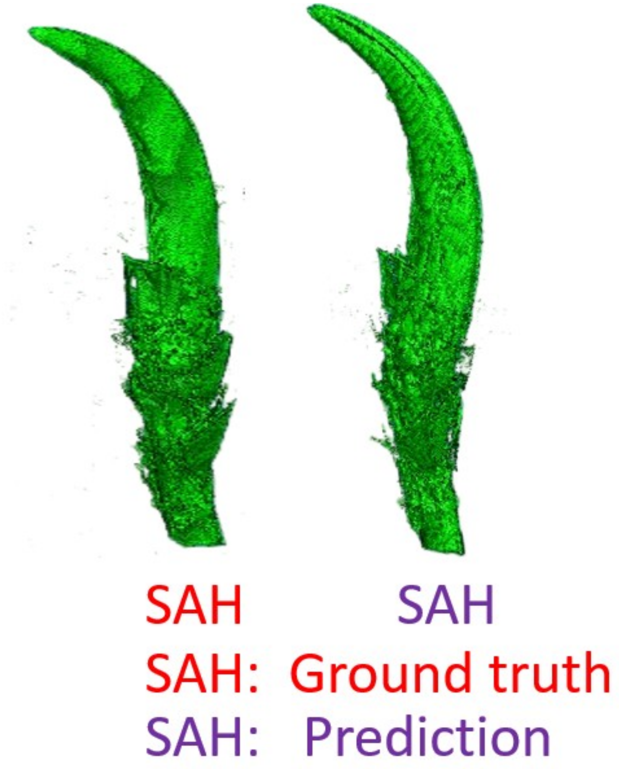


Figure B.12: Illustration of the qualitative comparison between the ground-truth and predicted 3D volume reconstructions of the hind claw of *Sceloporus arenicolus* (hind). The predicted volume was generated using the Inception-ResNet 2.5D model under the leave one out cross validation (LOOCV) framework

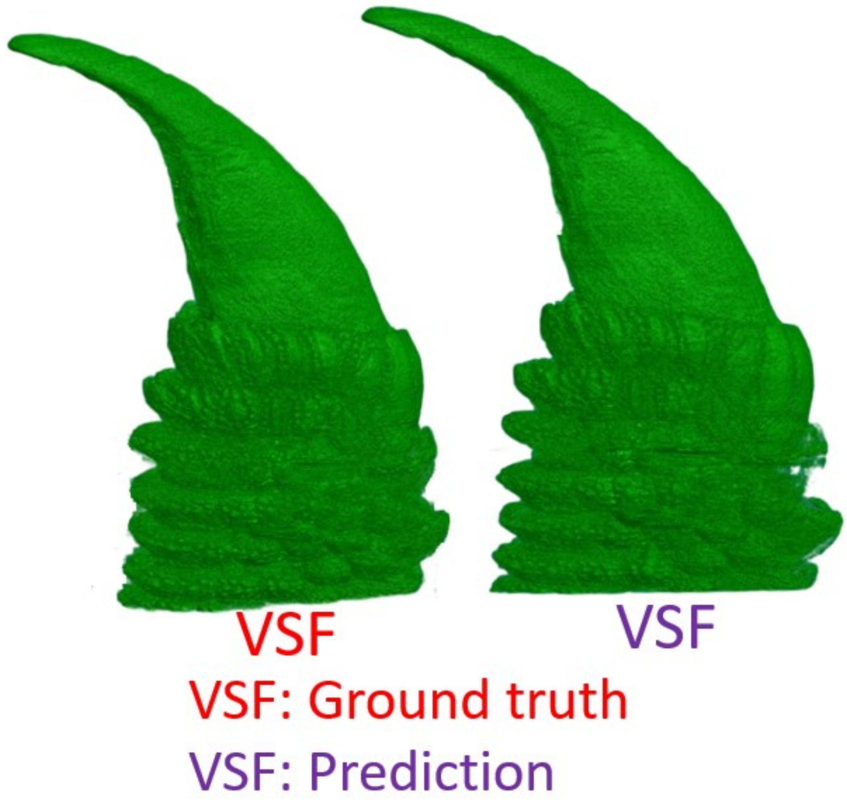


Figure B.13: Illustration of the qualitative comparison between the ground-truth and predicted 3D volume reconstructions of the hind claw of *Varanus salvator* (fore). The predicted volume was generated using the Inception-ResNet 2.5D model under the leave one out cross validation (LOOCV) framework

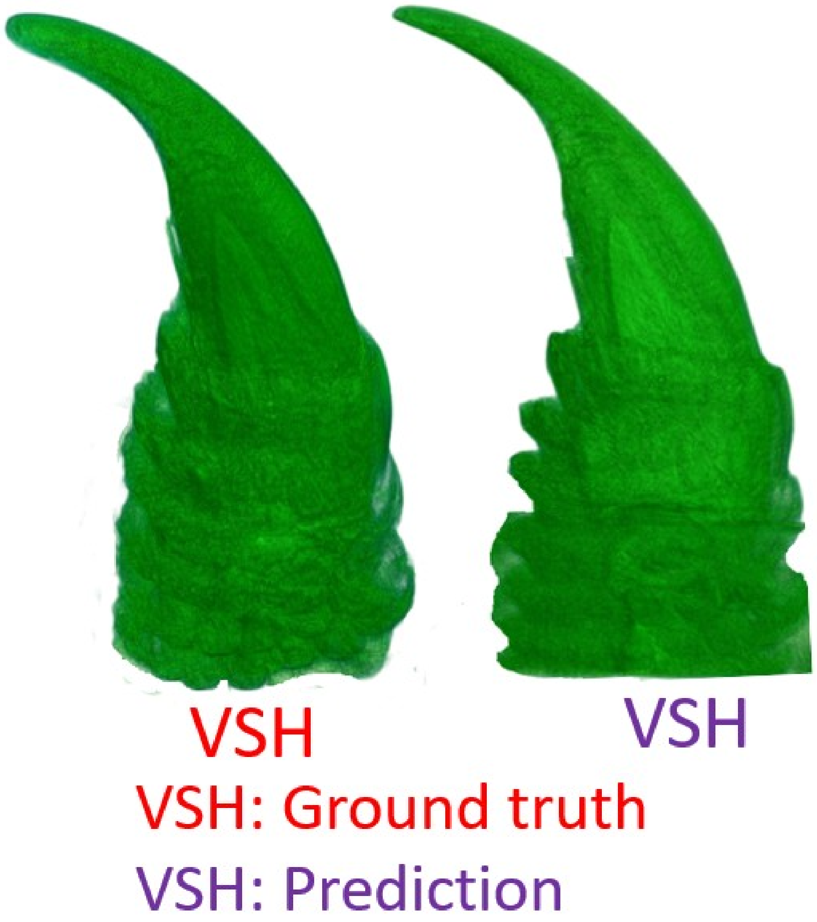


Figure B.14: Illustration of the qualitative comparison between the ground-truth and predicted 3D volume reconstructions of the hind claw of *Varanus salvator* (hind). The predicted volume was generated using the Inception-ResNet 2.5D model under the leave one out cross validation (LOOCV) framework
